# Consolidating *Ulva* functional genomics: Gene editing and new selection systems

**DOI:** 10.1101/2024.11.13.623382

**Authors:** Jonas Blomme, Júlia Arraiza Ribera, Olivier De Clerck, Thomas B. Jacobs

## Abstract

The green seaweed *Ulva compressa* is a promising model for functional biology. In addition to historical research on growth and development, -omics data and molecular tools for stable transformation are available. However, more efficient tools are needed to study gene function. Here, we expand the molecular toolkit for *Ulva*. We screened the survival of *Ulva* and its mutualistic bacteria on 14 selective agents and establish that Blasticidin deaminases (BSD or bsr) can be used as selectable markers to generate stable transgenic lines. We show that Cas9 and Cas12a RNPs are suitable for targeted mutagenesis and can generate genomic deletions up to 20 kb using the marker gene ADENINE PHOSPHORIBOSYLTRANSFERASE (APT). We demonstrate that targeted insertion of a selectable marker via homology-directed repair or co-editing with APT is possible for non- marker genes. We evaluated 31 vector configurations and found that bicistronic fusion of Cas9 to a resistance marker or incorporation of introns in Cas9 led to the most mutants. We used this to generate mutants in three non-marker genes using a co-editing strategy. This expanded molecular toolkit now enables us to reliably make gain- and loss-of-function mutants, additional optimizations will be necessary to allow for vector-based multiplex genome editing in *Ulva*.

## Introduction

The green seaweed *Ulva mutabilis/compressa* (*Ulva* hereafter) emerged as a suitable lab model organism since its original isolation from a beach in South Portugal by Bjørn Føyn in 1952 (Føyn, 1958). *Ulva* is a simple multicellular organism consisting of three cell types: blade, rhizoid and stem cells (Spoerner *et al*., 2012). Dedicated protocols for propagation and standardized growth medium have been developed, facilitating maintenance under laboratory conditions (Stratmann *et al*., 1996). Nuclear genome sequences of *Ulva* (De Clerck *et al*., 2018; Osorio *et al*., 2022) and a molecular toolkit for stable transformation are available (Oertel *et al*., 2015; Blomme *et al*., 2021). The ability to create gain- and loss-of-function mutants combined with genomic resources make *Ulva* a unique functional genomics model for green seaweed research (Blomme *et al*., 2023).

*Ulva* is resistant to several antibiotics and individuals can only undergo normal development with at least two mutualistic bacteria (Spoerner *et al*., 2012), limiting the development of additional transformation markers. At present, stable expression of a bleomycin resistance marker (BleR) allows for the selection of transgenic *Ulva* individuals (Oertel *et al*., 2015; Blomme *et al*., 2021). However, antibiotics like phleomycin (a type of bleomycin) cleave DNA and prolonged incubation could lead to the accumulation of mutations (Trastoy *et al*., 2005). Furthermore, reliance on one selection marker limits genetic engineering options, *e.g.* delivery of a second vector into a transformant. Multiple selectable markers have been described in other algae models such as *Chlamydomonas reinhardtti* (reviewed in (Doron *et al*., 2016; Perozeni & Baier, 2023)).

Mutant generation is essential to understanding gene function. One important feature of the original *Ulva* cultures was their “mutability”; spontaneous mutants with altered development (reviewed in (Wichard *et al*., 2015; Blomme *et al*., 2023)). Induced mutagenesis has also been achieved by UV-radiation (Fjeld, 1971; Løvlie, 1978) and random insertion of plasmid DNA (Oertel *et al*., 2015; Kwantes & Wichard, 2022). CRISPR/Cas has become the standard for targeted DNA mutagenesis in biology. CRISPR/Cas is a two-component system containing a guide RNA (gRNA) that directs a Cas nuclease to a specific DNA sequence. Upon target recognition, the nuclease generates a target-specific DNA double-strand break (Jinek *et al*., 2012). Mutations such as insertion or deletions (indels) then arise during error-prone DNA repair pathways such as Non-Homologous End Joining (NHEJ). Several alternatives to nucleases are also available for genome editing purposes, such as base editors that can modify nucleotides without a DNA double-strand break (Komor *et al*., 2016; Gaudelli *et al*., 2017).

While CRISPR/Cas is functional in different algal species, the reported efficiencies are low (<10%) and it is often necessary to target genes that confer a selectable phenotype upon mutation (Sproles *et al*., 2021). For example, a knock-out (KO) mutation in ADENINE PHOSPHORIBOSYLTRANSFERASE (APT) prevents the conversion of 2-fluoroadenine (2- FA) into the toxic 2-FluoroAMP. To date, CRISPR/Cas using APT mutagenesis has been described for three seaweed species: *Ectocarpus species 7*, *Saccharina japonica* and *Ulva prolifera* (Badis *et al*., 2021; Ichihara *et al*., 2021; Shen *et al*., 2023). *Ectocarpus* is the only example where additional genes were simultaneously co-edited with APT (Badis *et al*., 2021). CRISPR-mediated insertional mutagenesis in *U. prolifera* has also allowed for the tagging of the endogenous Rubisco with a fluorescent protein (Ichihara *et al*., 2024). In these reports, CRISPR components were delivered as ribonucleoproteins (RNPs). While useful, vector-based delivery systems have a number of advantages such as low cost and the versatility to use different CRISPR applications such as base editors. Efficient vector-based CRISPR will also allow the isolation of mutants without co-editing a selectable gene such as APT, as demonstrated in the red seaweed *Neopyropia yezoensis* (Wang *et al*., 2024). These reports clearly establish that CRISPR/Cas is a viable means to obtain targeted mutations in seaweeds, but more efficient tools are required to improve seaweed functional genomics.

In this report, we expanded the *Ulva* molecular toolkit by developing a novel Blasticidin resistance marker to select stable transformants. We also tested various CRISPR/Cas applications in *Ulva*: Cas9 and Cas12a nucleases, targeted insertion of template DNA and up to 20 kb genomic deletions, using both RNP and vector delivery. We used the APT marker to score genome editing efficiencies and screened 31 vector combinations for mutation efficiency. Our screen indicated that bicistronic fusion of Cas9 to BleR or incorporation of introns in Cas9 can increase APT mutagenesis. We also performed co-editing in four additional genes. We show that vector-based CRISPR mutagenesis is functional but relatively inefficient and that HDR-mediated insertion is a viable alternative. Further optimization will be necessary to allow for multiplex genome editing in *Ulva*.

## Material and methods

### Growth and cultivation

*Ulva mutabilis* Føyn “Wild-type” and “*slender*” strains are descendants of the original isolates from the South Atlantic coast of Portugal (Føyn, 1958). Both *Ulva* strains are maintained as haploid gametophytes and cultivated in a tissue culture chamber under long-day conditions (16-h light:8-h dark; 21°C; approx. 75 mE m^-2^ sec^-1^; Spectralux Plus NL-T8 36W/840/G13 fluorescent lamp) in standard Petri dishes (150x20 mm, SPL Life Sciences) containing 50-100 mL synthetic *Ulva* Culture Medium (UCM; (Stratmann *et al*., 1996; Boesger *et al*., 2018)). *Ulva* was grown and parthenogenetically propagated as previously described (Wichard & Oertel, 2010). All experiments in this report are generated using the *slender* strain unless stated otherwise.

For the sensitivity assay, UCM or Marine Broth (Sigma) were supplemented with 2- Fluoroadenine (2-FA; Sigma), 5-Fluoroorotic acid (5-FOA; Sigma), Blasticidin (Invivogen), Chloramphenicol (Sigma), Norflurazon (Sigma), G418 (Sigma), Hygromycin (Sigma), Oxyfluorfen (Sigma), Paromomycin (Sigma), Rapamycin (Sigma), Spectinomycin (Duchefa), Sulfometuron Methyl (SMM; Sigma) or Zeocin (Invitrogen) at the indicated concentration ranges. Chemical were dissolved in acetone (Oxyfluorfen, SMM), DMSO (2-FA, 5-FOA, Rapamycin, Sulfadiazin), ethanol (Norflurazon) or water (Blasticidin, Chloramphenicol, G418, Spectinomycin, Hygromycin, Paromomycin, Zeocin).

### Transformation

*Ulva* gametes were transformed using the PEG-mediated protocol previously described (Oertel *et al*., 2015; Boesger *et al*., 2018). For each transformation, 5 µg of DNA was used per plasmid. Plasmids were purified using a GeneJet Plasmid Miniprep kit according to manufacturer’s recommendations (Thermo Scientific). 50 µg/mL Phleomycin (Invivogen), 50 µg/mL Blasticidin or 10 µg/mL 2-FA was added 72 h after transformation. One additional volume of 40 mL selective medium was added on top after two weeks. Resistant germlings develop parthenogenetically from the transformed gametes. For each transformation experiment, no-vector controls were included to confirm proper selection. Escapes are sometimes observed (up to 4 individuals) in the no-vector empty controls with 2-FA selection.

For experiments where relative transformation or editing efficiency was scored, all plasmids and controls were transformed in the same transformation experiment (side-by-side). The number of gametes was counted before and after PEG-mediated transformation and all resistant individuals were counted to calculate relative transformation or editing efficiency for each transformation, summarized in **Table S1**.

### Genotyping

Genomic DNA was extracted from thalli of 1-month-old resistant individuals via a CTAB method (Clarke, 2009). Genotyping primers were designed to amplify a region of 700-900 bp around the CRISPR target sites. 10 µL PCR reactions were performed using the ALLin™ HS Red Taq Mastermix (HighQu). After amplification, 4 µL of PCR product was analyzed using gel electrophoresis and the remaining 6 µL was purified via magnetic beads (CleanNGS kit, CleanNA) as previously described (Blomme *et al*., 2022). PCR products were sent for Sanger sequencing (Mix2seq; Eurofins) and analysed using Geneious Prime software (Version 2024.0.7) and/or the ICE webtool (https://ice.synthego.com).

For the genotyping of large deletions, primers were designed around the targeted genome regions. To detect >1 kb deletions, a PCR was performed with the forward primer of the upstream target and the reverse primer of the downstream target. To confirm the quality of the extracted DNA, a control PCR was carried out to amplify an unrelated gDNA region (EFLa, UM025_0053) and all amplicons that corresponded to putative deletions were verified by Sanger sequencing (Mix2seq; Eurofins). All genotyping primers and gRNA sequences are listed in **Table S2**.

### RNP assembly

Purified crRNAs, trans-activating crRNA (tracrRNA), HiFi S.p. Cas9 Nuclease V3 and L.b. Cas12a Ultra were obtained from Integrated DNA Technologies (IDT) and assembled into RNP complexes following the manufacturer’s instructions with a Cas9:gRNA or Cas12a:crRNA ratio of 1:1 in NEBuffer 3.1 (NEB). 8µg of Cas9 or Cas12a complexed with equimolar ratios of gRNA/crRNA were transfected into *Ulva* unless stated otherwise.

For targeted insertion, BleR was cloned with 500bp genomic DNA upstream and downstream in pGG-AG-KmR for each target (**Table S3**). After assembly and verification, a fragment containing 50bp overhangs was amplified using Q5 polymerase (NEB). The amplicon was purified (DNA Clean & Concentrator-5; Zymo Research) and 1µg was included in the transformation with the RNP complex.

### Vector assembly

Vector assembly is based on the GreenGate cloning system and our existing molecular toolkit for *Ulva* (Lampropoulos *et al*., 2013; Blomme *et al*., 2021). Sequences for entry modules were obtained via gene synthesis (IDT, Neochromosome or TelesisBio) or PCR-amplification of the target DNA sequence using primers that contain a ∼20-bp overlap with the entry module using Q5 polymerase (NEB) or GXL (Takara Bio). After gel electrophoresis, the amplicons were gel purified (Zymoclean Gel DNA Recovery Kit; Zymo Research) and mixed with the predigested (BsaI; NEB) entry plasmid and NEBuilder® HiFi DNA Assembly Master Mix (NEB) for Gibson assembly. After assembly (15–60 min at 50°C), the cloning reactions were transformed into chemically competent DH5α *Escherichia coli* cells and plated on LB + 100 µg/mL Carbenicillin. PCR-mediated silent mutations were introduced to remove internal BsaI recognition sites. All *Ulva* gene and promoter sequences were cloned from genomic DNA, isolated via CTAB (Clarke, 2009) or Omniprep^TM^ for Plant (G-Biosciences).

The destination vector pGG-AG-KmR and its derivates contain BleR (Blomme *et al*., 2021). Similarly, the Blasticidin resistance cassettes (BlastR) were inserted in pGG-AG-KmR 3’ to the “G” BsaI recognition site. The BlastR cassettes were PCR amplified using pGG-AG- PIBT-KmR primers (Blomme *et al*., 2021) and gel purified. The pGG-AG-KmR was digested using EcoRV and NotI (Promega) and purified from gel. The BlastR amplicons were inserted in the digested pGG-AG-KmR vector via Gibson Assembly. In total, twelve destination vectors with different BlastR cassette configurations were generated (**Table S3**).

A two-step cloning procedure was used for the construction of modified destination vectors with B-E overhangs or to include up to six gRNA modules in a CRISPR vector (Decaestecker *et al*., 2019). First, a Golden Gate reaction was performed containing all desired fragments and two or three variable linkers containing AarI sites and the SacB counter selectable marker. The plasmids were digested with AarI and gel purified. The ccdB/cat cassette from pEN-L1-AG-L2 (Houbaert *et al*., 2018) was amplified using A-ccdB/cat-G or B-ccdB/cat-E primers (Blomme *et al*., 2021) and inserted into the digested vectors via Gibson Assembly. All destination vectors (**Table S3**) were transformed into One Shot ccdB Survival 2 T1^R^ Chemically Competent Cells (ThermoFisher).

For a Golden Gate reaction, 100 ng of each entry and destination vector were assembled in one reaction mix containing 10 U BsaI-HF-v2 (NEB), 200 U T4 Ligase (NEB), 1 mM ATP (Thermo Scientific), and 1x Cutsmart buffer (NEB). The Golden Gate reaction was run for 30 cycles (18°C [3 min] and 37°C [3 min]), followed by 5 min 50°C and 5 min 80°C. Approximately 2,5 µl of the assembly mix was transformed into DH5α *E. coli* cells and plated on LB + 25 µg/mL Kanamycin. Vectors were validated using Sanger sequencing (Mix2seq; Eurofins) or Oxford Nanopore whole-plasmid sequencing (Plasmidsaurus or Eurofins) and restriction digest, typically using NcoI (Promega), PstI (Promega), or XhoI (NEB). Vectors generated in this study (**Table S3**) are available at https://vectorvault.vib.be/ulva-mutabilis-toolkit.

### Confocal imaging

Thalli of 1-month-old resistant individuals were selected for imaging. Detailed imaging of cells was performed using an Olympus Fluoview (FV1000) confocal microscope with FluoView software (FV10-ASW 4.2). The following excitation and emission settings were used: YFP, 515 nm excitation with 530–545 nm emission and chlorophyll, 559 nm excitation with 650–750 nm emission. Laser intensity was adapted depending on expression variation between individuals. For each transgenic line, at least six independent individuals were screened and a representative image was selected.

## Results

### Developing a new antibiotic resistance marker

The development of *Ulva* antibiotic resistance makers is hampered by the requirement of two symbiotic bacteria for proper development (Spoerner *et al*., 2012) and there is anecdotal evidence of natural resistance to several selective agents (Oertel *et al*., 2015). We therefore performed a chemical screen to expand selection options. *Ulva* germlings were transferred to a series of selective or control media and growth was evaluated after two weeks. We used 14 different selective agents: 2-Fluoroadenine (2-FA), 5-Fluoro-orotic acid (5-FOA), Blasticidin, Chloramphenicol, G418, Hygromycin, Norflurazon, Oxyfluorfen, Paromomycin, Rapamycin, Spectinomycin, Sulfadiazin, Sulfometuron Methyl (SMM) and Zeocin. These chemicals represent a broad range of antibiotics and herbicides and are used with other (micro)algae such as *Chlamydomonas* (reviewed in (Doron *et al*., 2016)). *Ulva* growth was unaffected by the tested concentrations of G418, Hygromycin, Oxyfluorfen, Paromomycin, Rapamycin and Zeocin. Spectinomycin and Sulfadiazin did not affect *Ulva* survival, but did inhibit growth at higher concentrations. Six chemicals had a negative effect on *Ulva* survival at higher concentrations: 2-FA, 5-FAO, Blasticidin, Chloramphenicol, Norflurazon and SMM (**Fig. 1a; Fig. S1**).

**Figure 1.**
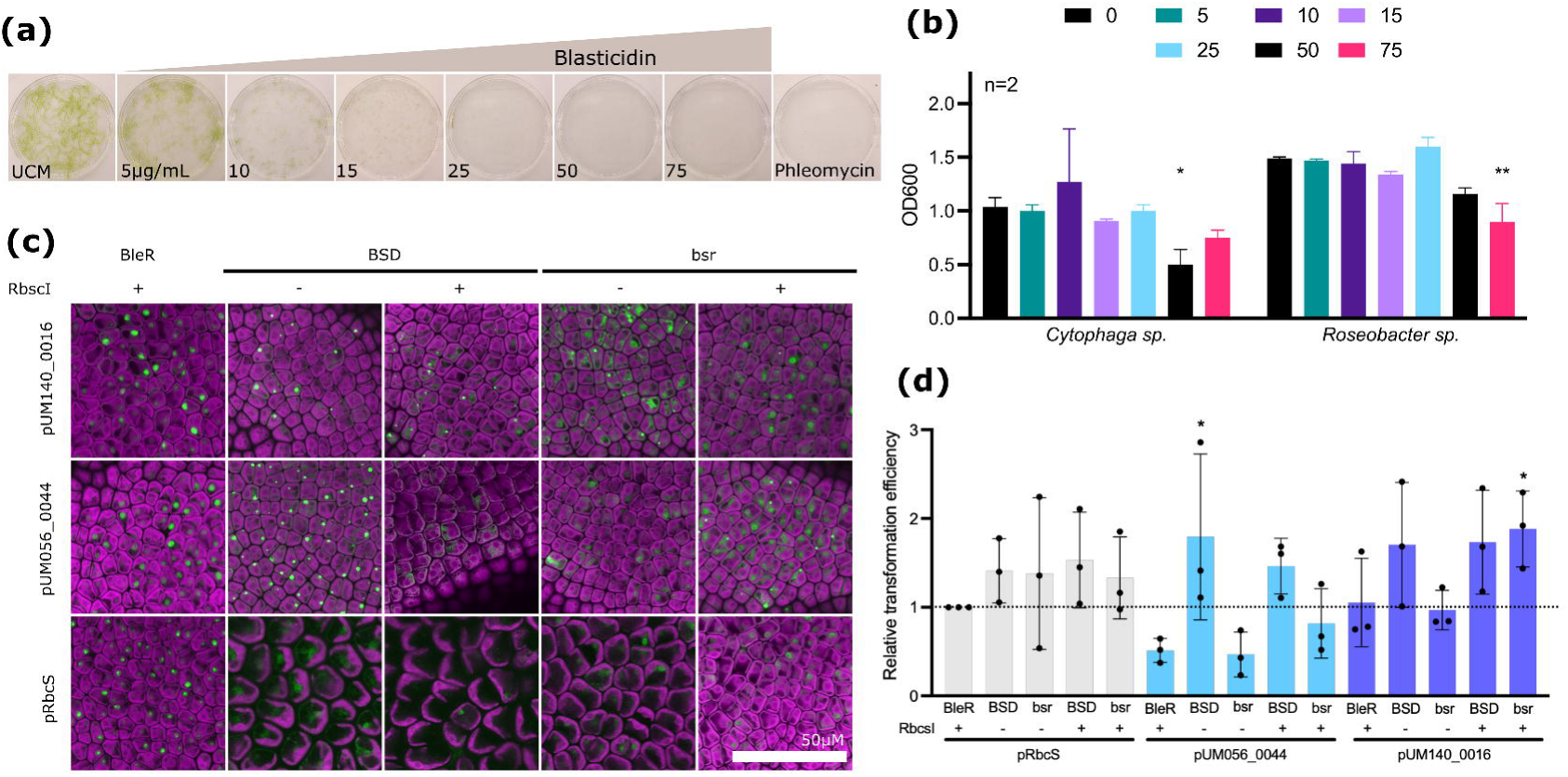
A new antibiotic resistance marker for *Ulva*. (a) Sensitivity assay for Blasticidin, including no treatment (UCM) and Phleomycin (50µg/mL) controls. *Ulva* germlings were transferred to increasing concentrations Blasticidin. Cultures were photographed two weeks after transfer. All values represent concentrations in µg/mL. (b) Sensitivity assay for Blasticidin. *Cythopaga sp.* MS6 and *Roseobacter sp.* MS2 were inoculated at increasing concentrations Blasticidin (0-75 µg/mL). All values represent average OD600 (±SD; n=2) two weeks after inoculation. Significant differences from the blank control are indicated (ANOVA with *post hoc* Dunnett multiple comparison test; * = P >0,05 and ** = P >0,01). (c) Overview of transgene expression. BSD or bsr were directly fused to YFP under control of one of three promoters (pUM140_0016, pUM056_0044 or pRbcS) and a RbcsI sequence (+) or not (-). A BleR-YFP fusion is included as control. Magenta: Chlorophyll autofluorescence; Green: YFP signal. Scale = 50µM. (d) Relative *Ulva* transformation efficiency with BSD or bsr under control of one of three promoters (pUM140_0016, pUM056_0044 or pRbcS) and a RbcsI sequence (+) or not (-). Values represent relative transformation efficiency compared to pRbcS-RbcsI- BleR-tRbcS ±SD (n=3). Significant differences from the control are indicated (ANOVA with *post hoc* Dunnett multiple comparison test; * = P >0,05).

To determine which selective agents were compatible with symbiotic bacterial growth, *Cytophaga sp*. MS6 and *Roseobacter sp.* MS2 were transferred to liquid growth medium containing the same concentration range of chemicals and the optical density (OD600) was measured to assess growth. Both bacteria were highly sensitive to 5-FAO and Chloramphenicol after two weeks. Growth was also slower on higher concentrations of G418, Hygromycin and Sulfadiazin (**Fig. 1b; Fig. S2**).

These results pointed to Blasticidin as a candidate selective agent as *Ulva* is sensitive at low concentrations while bacterial growth is not affected. Blasticidin inhibits protein synthesis and resistance can be conferred by expressing either of two Blasticidin-S deaminases: BSD, originally isolated from *Aspergillus terreus* (Kimura *et al*., 1994), or bsr from *Bacillus cereus* (Izumi *et al*., 1991). We cloned both genes and translationally fused them to an N-terminal YELLOW FLUORESCENT PROTEIN (YFP). We tested three previously characterized promoters to control expression (pRbcS, pUM140_0016 or pUM056_044; (Oertel *et al*., 2015; Blomme *et al*., 2021)), with or without the first intron of UM023_0100 encoding the small subunit of Rubisco (RbcsI; (Oertel *et al*., 2015; Blomme *et al*., 2021)) upstream of the start codon. Both BSD-YFP and bsr-YFP fusion proteins were expressed in individuals selected on Blasticidin and localized to the nucleus and cytoplasm as expected (**Fig. 1c**). All resistance markers conferred Blasticidin resistance, but BSD in combination with pUM056_0044 without RbcsI and bsr under control of pUM140_0016 with RbcsI resulted in significantly more resistant individuals compared to the pRbcS-RbcsI-BleR-tRbcS control (P > 0,05; ANOVA with *post hoc* Dunnett multiple comparison test; **Fig. 1d; Table S1**). All resistance markers were integrated into pGG-AG-KmR to create 12 new destination vectors that complement our existing vector toolkit ((Blomme *et al*., 2021); **Fig. S3; Table S3**). To facilitate expression vector assembly, we designed four destination vectors where only the gene of interest and tag need to be inserted (*e.g.* pGG-pUM140_0016-BE-NOST) for all pRbcS constructs driving BSD or bsr without YFP (**Fig. S3; Table S3**). Thus, we have a new selectable marker system for *Ulva*.

### *Ulva* vector optimization for CRISPR mutagenesis

Complementary to our efforts to develop tools to generate stable overexpression lines (Blomme *et al*., 2021), we initiated several experiments to establish CRISPR/Cas vectors for *Ulva.* Two putative Pol III promoter elements were isolated upstream of U6 genes UM003_0070 (pUMU6-1) and UM026_0006 (pUMU6-2) and cloned with the SpCas9 scaffold and two BbsI restriction enzyme sites to insert gRNA spacer sequences in a one-step cloning reaction ((Decaestecker *et al*., 2019); **Table S3**). We synthesized *Chlamydomonas* and *Ulva* codon-optimised Cas9 (CrCas9 and UmCas9, respectively) sequences and coupled them to previously generated promoters or RbcsI variants (**Table S3;** (Blomme *et al*., 2021)). Approximately 5 µg of each plasmid was transfected into gametes and selected on phleomycin for 4 weeks. In an initial round of experiments, we tested a variety of vector configurations including different promoters, inclusion of an intron sequence, codon optimizations, protein fusions and nuclear localization signals (NLSs). We selected two target genes: URA3 (UM011_0242) which should generate auxotrophic mutants and CCM1 (UM072_0060) which encodes a transcription factor with a putative role in carbon capture (**Fig. S4**). In total, we tested 44 different vector configurations and genotyped 424 phleomycin-resistant individuals, but no mutations were observed (**Table S1**).

To check whether our novel Cas9 coding sequences were competent to perform genome editing, CrCas9 and UmCas9 codon optimized versions were transformed into *Arabidopsis thaliana* Col-0 targeting PHYTOENE DESATURASE3 (PDS3) and GLABRA1 (GL1). Both coding sequences led to similar numbers of mutants as the Arabidopsis codon-optimized Cas9 in the T1 generation, confirming that CrCas9 and UmCas9 were indeed functional (**Fig. S5**).

### RNP-based genome editing in *Ulva*

Given our inability to induce mutations using various vector configurations, we decided to try RNPs targeting the APT marker gene as mutants could be positively selected via 2-FA (**Fig. 2a,b; Fig. S1**). Three targets were designed in the second exon of APT (UM050_0062; **Fig. S4**). To test if the RNP complexes could at least function on naked DNA, we performed an *in vitro* digest with RNPs targeting URA3, CCM1 and APT. Seven out of eight regions were clearly cleaved, indicating their functionality *in vitro* (**Fig. S6**).

**Figure 2.**
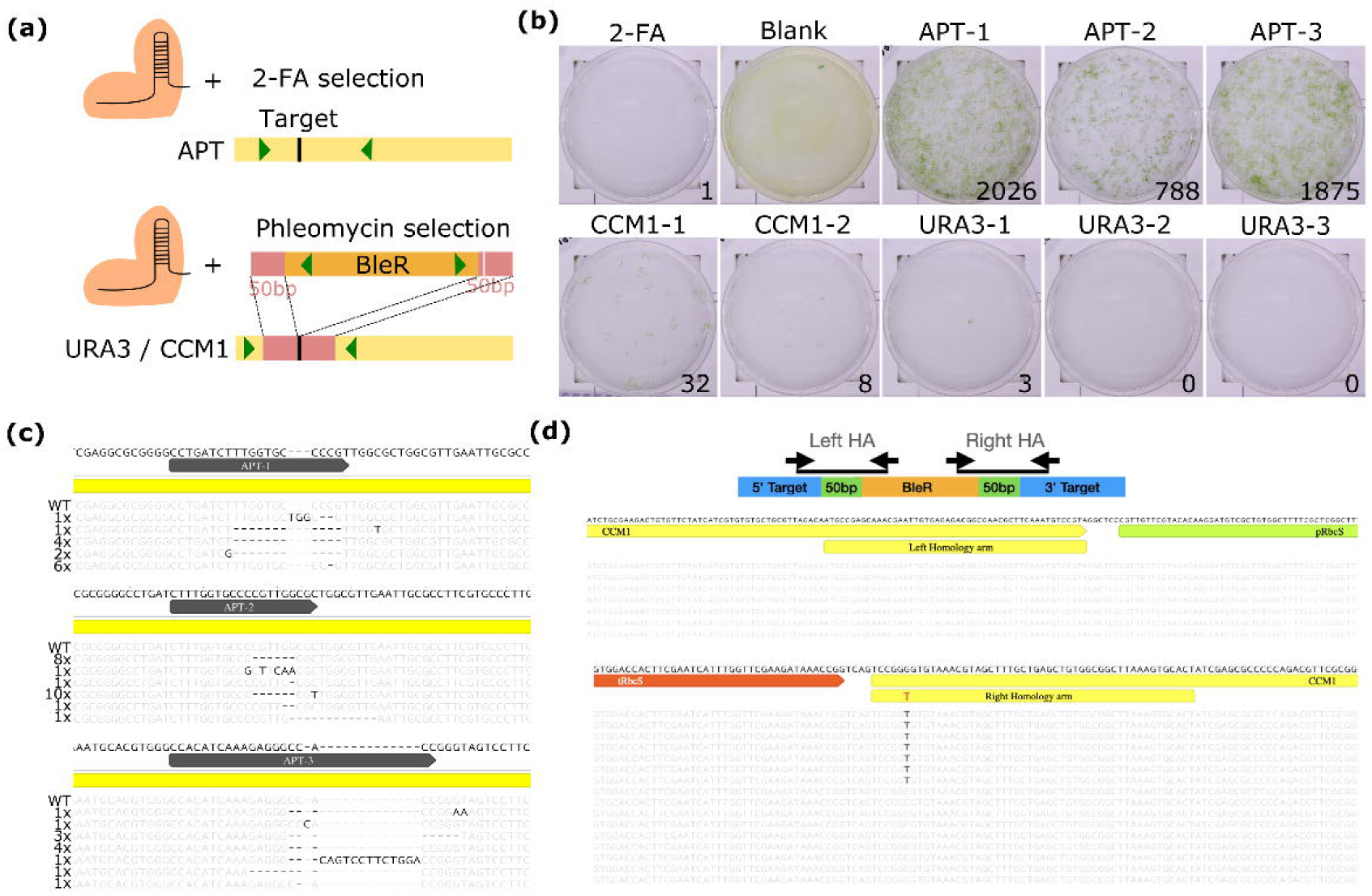
RNP-based genome editing in *Ulva*. (a) Overview of experiment. Targeted genome editing of the APT marker gene using a Cas9 RNP, resistant individuals were selected by adding 2-FA to the transfected cells. Non-marker genes such as URA3 and CCM1 can be targeted with a Cas9 RNP and a repair template, a dsDNA fragment containing the BleR cassette and 50bp overhang (red) to the target site (black). A SNP was introduced in the 3’ overhang to destroy the PAM site (grey). Selection using Phleomycin is possible after targeted insertion. Green arrowheads indicate genotyping primers. (b) Result of RNP transfection and 2-FA (APT-1, APT-2 and APT-3) or Phleomycin (CCM1-1, CCM1-2, URA3-1, URA3-2 and URA3-3) selection. Number of resistant individuals is indicated on the transfection plates. (c) Observed indels in 2-FA resistant individuals. Sanger sequencing reads were aligned to the reference gene sequence using Geneious software. The target site (APT-1, APT-2 and APT-3) is annotated in grey on the coding sequence (yellow). Every mutant read contains the number of genotyped individuals with the same indel pattern. An untransformed control (WT) is included. (d) Targeted insertion of BleR in CCM1 and URA3. PCR strategy to amplify the left or right homology arm (HA) is shown. Below are Sanger sequencing reads for CCM1-1 target at left and right border, visualized using Geneious software. Each lane represents one phleomycin-resistant individual. One SNP in the right HA is indicated in red. See **Fig. S11** for additional target sites.

The RNP complexes were transfected into *Ulva* gametes and individuals were selected using 10 µg/mL 2-FA. In total, there were 2028 2-FA resistant individuals for APT-1, 788 for APT- 2 and 1875 for APT-3, which amounts to an efficiency of 7 E^-05^- 2 E^-04^. Several 2-FA resistant individuals (14 for APT-1, 22 for APT-2 and 17 for APT-3) were genotyped by Sanger sequencing to confirm 2-FA resistance was caused by mutations in APT. All 2-FA resistant individuals contained indels at the target sites. Different repair outcomes were identified, including 1-11bp deletions, 1-13bp insertions and SNPs (**Fig. 2c**) and we obtained similar results in an independent *Ulva* strain called Wild-type (**Fig. S7**). Reducing the amount of Cas9 protein, from 8 µg to 2-6 µg Cas9 still resulted in hundreds of 2-FA resistant individuals (**Fig. S8**). We also tested LbCas12a RNPs by targeting APT. We designed two crRNAs targeting similar regions as the Cas9 APT-1 and APT-3 targets. Similar to the Cas9 experiment, Cas12a RNPs induced indels in all genotyped individuals (17 for APT-1, and 23 for APT-2; **Fig. S9**). We observed 43x more resistant individuals when APT-1 was targeted compared to APT-2, illustrating a clear difference in crRNA efficiency. These results indicated that RNP based mutagenesis of the APT marker gene is straightforward in *Ulva*.

The ability to create targeted edits and our previously described vector tools (Blomme *et al*., 2021) should allow for genetic complementation, a fundamental test to restore a wild-type phenotype in a mutant. To test this, 2-FA resistant individuals were transformed with a complementation construct containing an APT-YFP fusion whose expression was controlled by the strong pUM140_0016 promoter (pUM140_0016-APT-YFP-NOST) for all three APT targets. Pools of T0 events were selected on phleomycin and propagated to the next generation. 24 T1 individuals were transferred to UCM medium with or without 2-FA. All individuals in 2-FA medium died or had severely disturbed growth, indicating that the APT- YFP fusion protein is functional and can convert 2-FA into the toxic 2-FluoroAMP (**Fig. S10a**). Two T1 individuals per complemented line were genotyped by Sanger sequencing. Primers were designed to capture both the endogenous, mutated APT alleles and the complementation construct. Multiple alleles were observed, including the wild-type allele in different proportions, suggesting that multiple copies of vector DNA were integrated (**Fig. S10b**). These results show that we can complement gene function in *Ulva* and APT provides us with an additional transgenic selectable marker system.

The efficiency of APT mutagenesis using RNPs indicated that *Ulva* is competent for CRISPR-based DNA cleavage. To try to make KO mutations in other target genes, we tested targeted insertion using RNPs and a DNA repair template containing a BleR cassette (1805 bp) with 50 bp homology arms for homology-directed repair (HDR; **Fig. 2a**). Phleomycin resistant individuals were obtained for three of the five targets (CCM1-1, CCM1-2, and URA3-1; efficiencies 3 E^-07^ – 3 E^-06^; **Fig. 2b**) and harvested for genotyping. To determine if resistance was due to targeted integration or random insertion, primers were designed to amplify both the target site and either the left or right homology arms of BleR (**Fig. 2d**). Left and right borders were amplified in 14 out of 32 individuals for CCM-1, 1 out of 8 for CCM1-2 and none of the 3 individuals for URA3-3 (**Fig. S11**). The amplicons were Sanger sequenced and perfect insertions were confirmed in 25 out of 28 reads. The exceptions being a 1 bp deletion in two CCM1-2 right border reads and one 17 bp deletion in the terminator of BleR in a CCM1-1 right border read. Interestingly, a SNP included in the PAM sequence of the CCM1-1 repair template was incorporated in only 7/18 individuals (**Fig. 2d; Fig. S11**). These results demonstrate that targeted insertion of BleR via HDR is viable for gene KO in *Ulva*.

### RNP editing of two targets

Given that we could easily recover hundreds to thousands of 2-FA resistant individuals when targeting a single APT site, we next tested if two targets can be edited at the same time. We started by simply targeting two regions less than 200 bp apart within the *APT* gene using Cas9 or LbCas12a RNPs. For Cas9, 41 2-FA resistant individuals were genotyped by Sanger sequencing, 21 contained a deletion of the genomic region between APT-1 and APT-3 (161 bp), 4 individuals had an inversion and the remaining 16 individuals had smaller indels at one (8) or both (8) target sites. For LbCas12a, 3/24 had a deletion of the DNA between both targets (136 bp) and the remaining 21 individuals had indels at APT-1 and/or APT-2 (**Fig. 3a,b; Fig. S9**).

**Figure 3.**
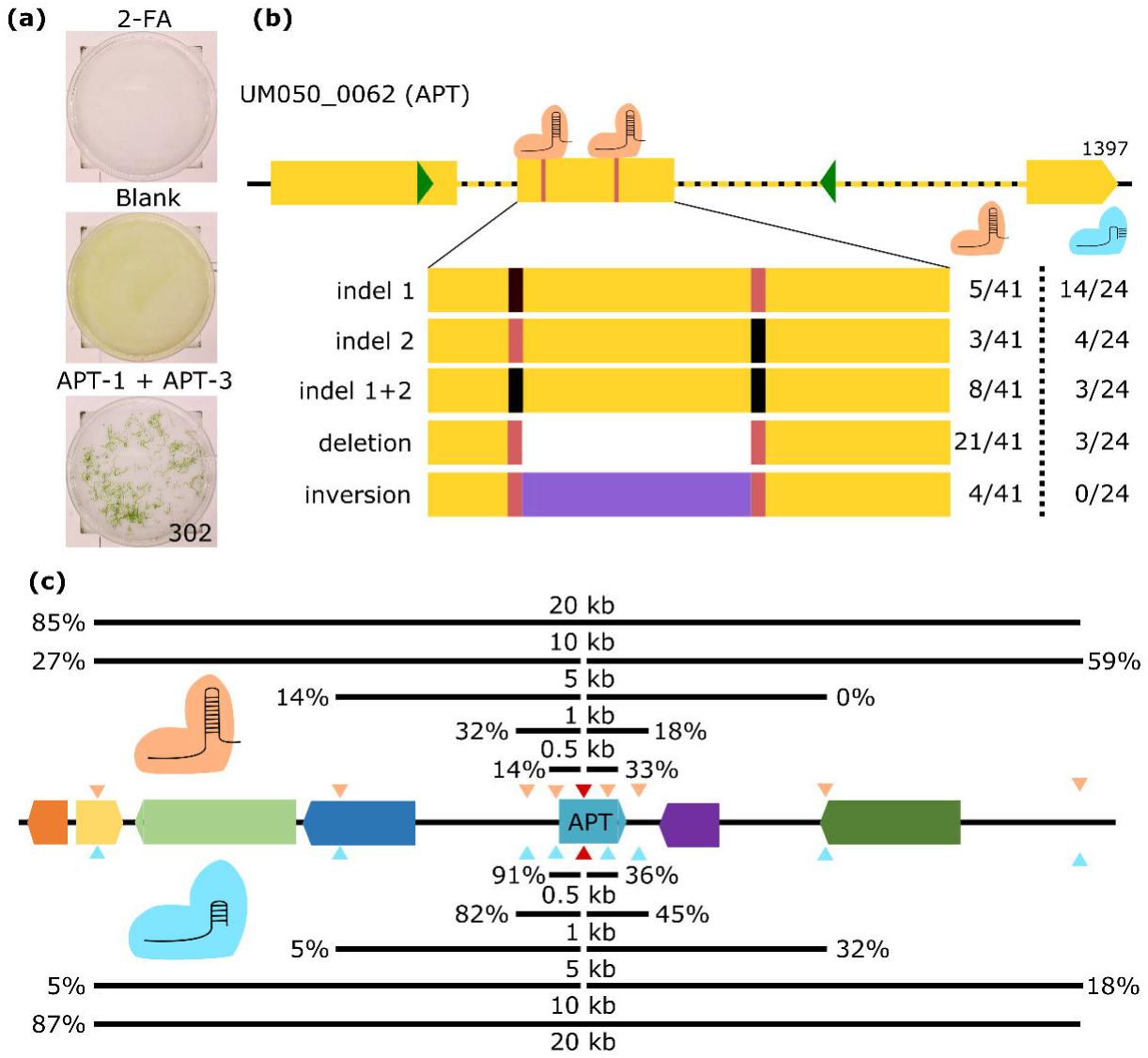
RNP editing of two targets simultaneously in *Ulva*. (a) Result of APT-1 and APT-3 RNP co-transfection and 2-FA selection. Number of resistant individuals is indicated on the transfection plate. (b) Repair outcomes of APT-1 and APT-3 RNP co-transfection in 2-FA resistant individuals. Both target sites (red) are localized in the second exon of APT. Genotyping primer position is indicated with green arrowheads. Experiment was done using Cas9 (orange) or LbCas12a (blue) nuclease. Sanger sequencing reads were aligned to the reference gene sequence and summarized according to one of five outcomes: indels at either or both targets, a deletion of the DNA between both target sites and an inversion of the DNA region between both sites (purple). The number of identified individuals is listed for each outcome and nuclease. (c) Large deletion experiment. Targets were designed at 0.5, 1, 5 or 10kB distance of APT-1, both up- and downstream and for Cas9 (orange) and LbCas12a (blue) nucleases. The target sites are represented on a schematic overview of the genetic region with arrowheads. In each experiment APT-1 (red arrowhead) and a second target were co-transfected, the percentage of 2-FA resistant individuals with a deletion identified by PCR is indicated for each combination. All results of Cas9 RNPs are visualized above the overview of the genetic region, results of LbCas12a RNPs are shown below. See also **Table S4**.

Motivated by the observation that both nucleases created relatively large deletions, and Cas9 did so in approximately half of the events, we tested if even larger chromosomal regions could be as easily removed. We targeted both APT-1 and a second target at 0.5, 1, 5 or 10 kb distances, both up- and down-stream of APT-1 using both nucleases. We also included RNPs targeting the two most distal loci to create a 20 kb deletion. This region contains five additional protein-coding genes (**Fig. 3c**). For each distance and nuclease, 20-23 individuals were genotyped for deletions via PCR and gel electrophoresis. We readily detected large deletions for almost all combinations, with 5-91% of genotyped individuals containing a deletion, which was verified for most individuals by Sanger sequencing (**Fig. 3c; Table S4**). There was only one combination, Cas9 APT-1 and 5 kb downstream, where no deletions were observed, suggesting a less efficient gRNA. Most surprisingly, ∼85% of the resistant individuals from both the Cas9 and Cas12a 20 kb deletion treatments contained large deletions (**Fig. 3c; Table S4**). Thus, kilobase-scale deletions can be readily obtained in *Ulva* in and around the *APT* target gene.

The KO of the *APT* gene provided an excellent selectable marker and the large deletion experiments demonstrated that we could efficiently produce multiple DNA breaks. Therefore, we wondered if we could use a co-editing strategy (Badis *et al*., 2021) to generate KO lines. In co-editing experiments, mutagenesis at one locus is positively correlated with mutations at another locus. At least two gRNAs are delivered to the cells; one in a gene of interest and a second to target a selectable marker (*e.g.* APT). We tested the co-editing strategy by co-transfecting APT-1 and CCM1-1 RNPs and selected 2-FA resistant individuals. As expected, all 46 genotyped individuals had indels at the APT-1 target site. However, the CCM1-1 target was not edited in these nor in additional individuals (78 in total). We also tested other targets by co-transfecting the APT-1 RNP with four additional RNPs. For each co-editing transfection, 71-88 2-FA resistant individuals (380 in total) were genotyped by Sanger sequencing. Of these, only one individual was mutated at the second target site (CCM1-2) and it contained both wild-type and mutated alleles (**Fig. S12**). Thus, co-editing can produce mutated lines, but the efficiency is very low.

### Simplex vector-based Cas9 editing

Our next approach was to develop a vector-based approach as it is more flexible (*e.g.* base editors) and scalable (multiplex CRISPR) than RNP-mediated mutagenesis. We incorporated the APT-1 gRNA into some of the vector configurations previously generated (**Fig. 4a; Table S1**). Both promoter and NLS have been identified as key components to optimize CRISPR efficiency (Wang *et al*., 2015; Feng *et al*., 2018; Castel *et al*., 2019; Grützner *et al*., 2021; Gaillochet *et al*., 2023; Develtere *et al*., 2024). We coupled UmCas9 to the strong pUM140_0016 promoter and incorporated either a SV40 or N7 NLS with a direct or bicistronic (F2A) fusion to YFP. The incorporation of introns has been shown to enhance transgene expression in algae and Cas9 expression in plants (Lumbreras *et al*., 1998; Perozeni *et al*., 2020; Grützner *et al*., 2021; Amendola *et al*., 2023) and was essential for transgene expression in *Ulva* (Oertel *et al*., 2015; Blomme *et al*., 2021). We therefore inserted RbcsI either upstream of the UmCas9 transgene or incorporated it 1, 2, 4 or 10 times in the CDS using intronserter (Jaeger *et al*., 2019). We also coupled UmCas9 expression to the BleR selectable marker via a bicistronic (F2A, P2A and T2A) or direct fusion under control of pRbcS, a strategy also employed in *Neopyropia* (Wang *et al*., 2024). We generated all variants with both *Ulva* U6 promoters, 30 vector configurations in total (**Fig. 4a; Table S1**).

**Figure 4.**
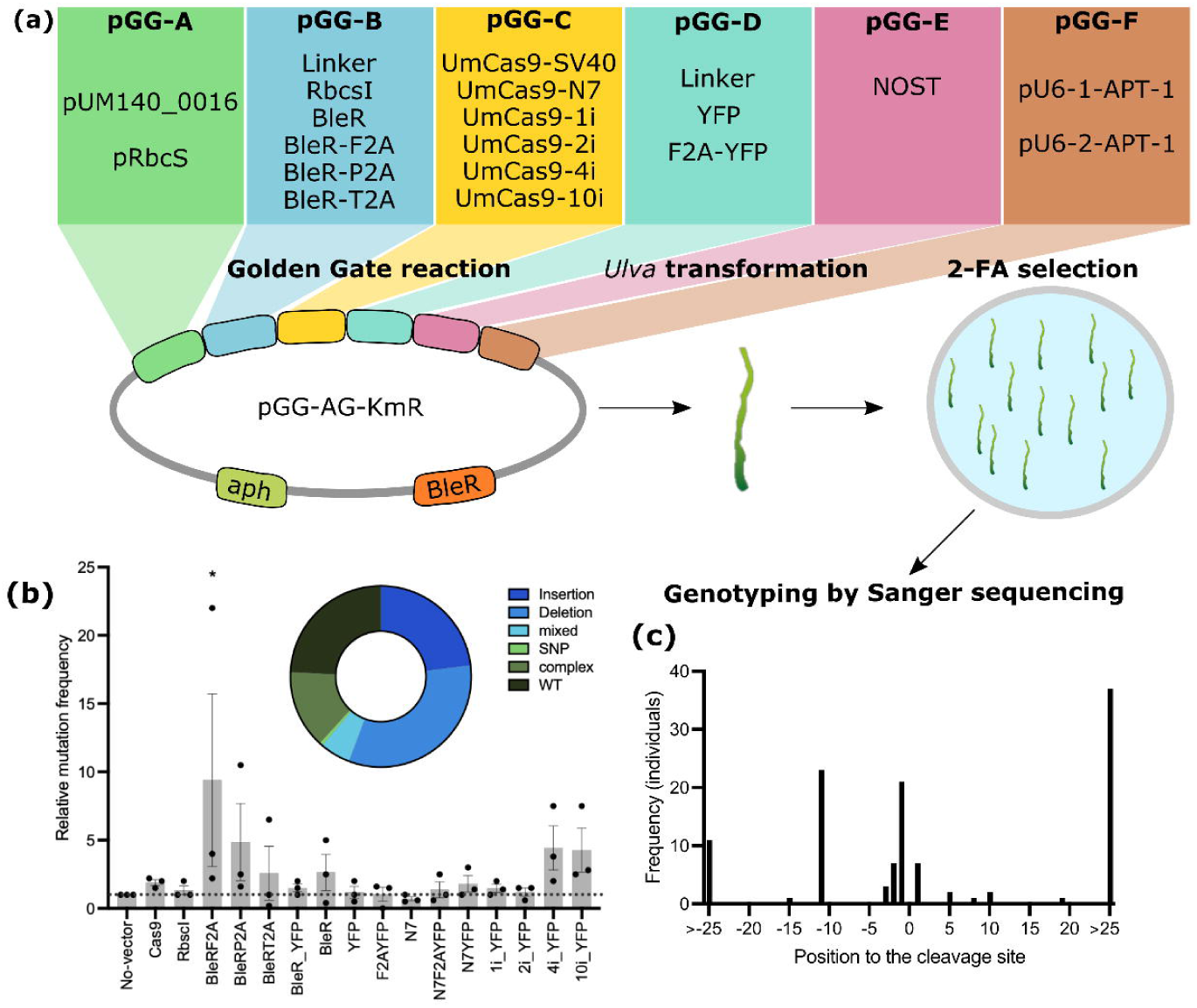
Vector-based genome editing in *Ulva* (a) Vector screen set-up for *Ulva* genome editing. Different entry modules (pGG-A to pGG-F) are depicted, thirty different combinations were assembled via Golden Gate cloning in an expression vector. The vectors were transformed in *Ulva*, after one month of selection the number of 2-FA resistant individuals was counted and genotyped by Sanger sequencing. (b) Relative mutation frequencies of 16 vector variants and a no-vector control. 2-FA resistant individuals were counted and this value was put relative to the empty no-vector control per independent experiment (n=3, values are average ±SE). Significant differences from the blank control are indicated (ANOVA with *post hoc* Dunnett multiple comparison test; * = P >0,05). See also **Table S1**. Insert: summary of mutation type based on Sanger sequencing results (n=165). (c) Frequency distribution of CRISPR-induced insertions and deletions in *Ulva*. The number of individuals per insertion or deletion at each position relative to the target site is shown, individuals with insertions or deletion >25bp were binned. n=116.

Given the large number of vector architectures, we selected 12 with two expression cassettes for an initial test (**Table S1**). After *Ulva* transformation and 2-FA selection, 1-23 resistant individuals were obtained per vector. Genotyping by Sanger sequencing confirmed mutations in APT, including small indels and larger insertions of vector DNA up to 343 bp (**Table S1**). A few individuals were not edited at the *APT* target site in six vector variations and are probably escapes from selection. With a functional CRISPR vector system in hand, we screened all thirty vectors targeting APT-1. After transformation and 2-FA selection, 1-82 resistant individuals were obtained across all vectors (mutation frequency 6 E^-07^ – 5 E^-05^).

Overall, the number of resistant individuals using the U6-1 promoter is ∼6x higher compared to vector configurations with the U6-2 promoter (**Table S1**; mean mutation frequency 1,2 E^-05^ versus 2,4 E^-06^; n=15; P>0,01; Two-tailed t-test). The vectors with the highest frequency of resistant individuals included a BleR-F2A-Cas9 bicistronic construct and Cas9 containing four RbcsI sequences (**Table S1**).

Since we observed a relatively large difference in number of resistant individuals, we aimed to test if one vector configuration consistently results in more mutant individuals and which mutations are generated. To test this, we transformed all 16 vectors containing U6-1 in triplicate in *Ulva*, selected on 2-FA, and quantified and genotyped the resistant individuals. This screen again resulted in a large variation of resistant individuals (0-63; mutation frequency 7 E^-08^ – 3 E^-06^). Only the BleR-F2A-Cas9 bicistronic construct resulted in significantly more resistant individuals (∼9x) compared to the no-vector control (P>0,05; ANOVA with *post hoc* Dunnett multiple comparison test; **Fig. 4b**), illustrating that we are at the limit of detection. Vectors containing BleR-P2A-Cas9 and Cas9 containing four or ten RbcsI sequences produced 4-5x more resistant individuals compared to the no-vector control. We harvested all resistant individuals to investigate the induced mutations at the target site. We were able to obtain high-quality Sanger sequencing data for 170 out of 326 individuals. We observed edits at the target site for all vector configurations, except for untagged UmCas9 with an N7 NLS (N7), but we also observed 1-3 individuals without mutations in the spacer for all vectors (**Fig. 4b; Table S1**). The most common repair outcomes were insertions (+1 bp to +349 bp) or deletions (-1 bp to -195 bp), but also SNPs and more complex edits were observed, usually the presence of >1 allele or a combination of indel and SNP (**Fig. 4b,c**). The >20 bp insertions mapped at least partly to vector DNA in 29 out of 37 individuals. Interestingly, two different 11 bp deletions were a frequent DNA repair outcome, both of which are predicted by inDelphi (Shen *et al*., 2018) and are associated with a microhomology of 2 or 4 bp. Taken together, vector-based genome editing using Cas9 is functional but inefficient in *Ulva*. We identify a few vector configurations that increase mutation frequency of the APT marker gene and could be the basis for additional optimization.

### Duplex vector-based Cas9 editing

While the number of 2-FA resistant individuals was orders of magnitude lower when using vectors as compared to RNPs, we still tested if our current vectors could mutate two genes simultaneously. The expression vector containing Cas9 with four introns and YFP was selected. We used a co-editing strategy, co-transforming APT-1 with APT-3, CCM1-1, CCM-2, URA3-1, URA3-2, URA3-3 or six additional gRNAs targeting 12-kD FK506 BINDING PROTEIN (FKB12; UM003_0238) and two genes where DNA insertion lines have been described (UM005_0337 and UM033_0004; (Kwantes & Wichard, 2022); **Fig. S4**). The APT-1 gRNA was expressed by U6-1 or U6-2 promoters and the second target was expressed with the stronger U6-1 (**Fig. 5a**). After transformation and selection in 2-FA, 4 to 16 resistant individuals per transformation were genotyped. Similar to the RNP co-editing strategy, we readily obtained mutants at APT-1 and/or APT-3 target sites (mutation frequency 4 E^-06^ – 1 E^-04^). We also recovered mutants in four other target sites: FKB12-2 (1/12 individuals), UM005_0337-1 (5/13 individuals), UM005_0337-2 (1/4 individuals) and UM033_0004-1 (2/19 individuals; **Fig. 5b**). The DNA repair products were smaller indels and vector DNA insertions up to 169 bp. These results show that co-editing was successful in 4/11 non-APT target sites at reasonable efficiencies (8 – 38% of genotyped individuals) and suggest that the URA3 and CCM1 targets are less functional compared to those for FKB12, UM005_0337 and UM033_0004.

**Figure 5.**
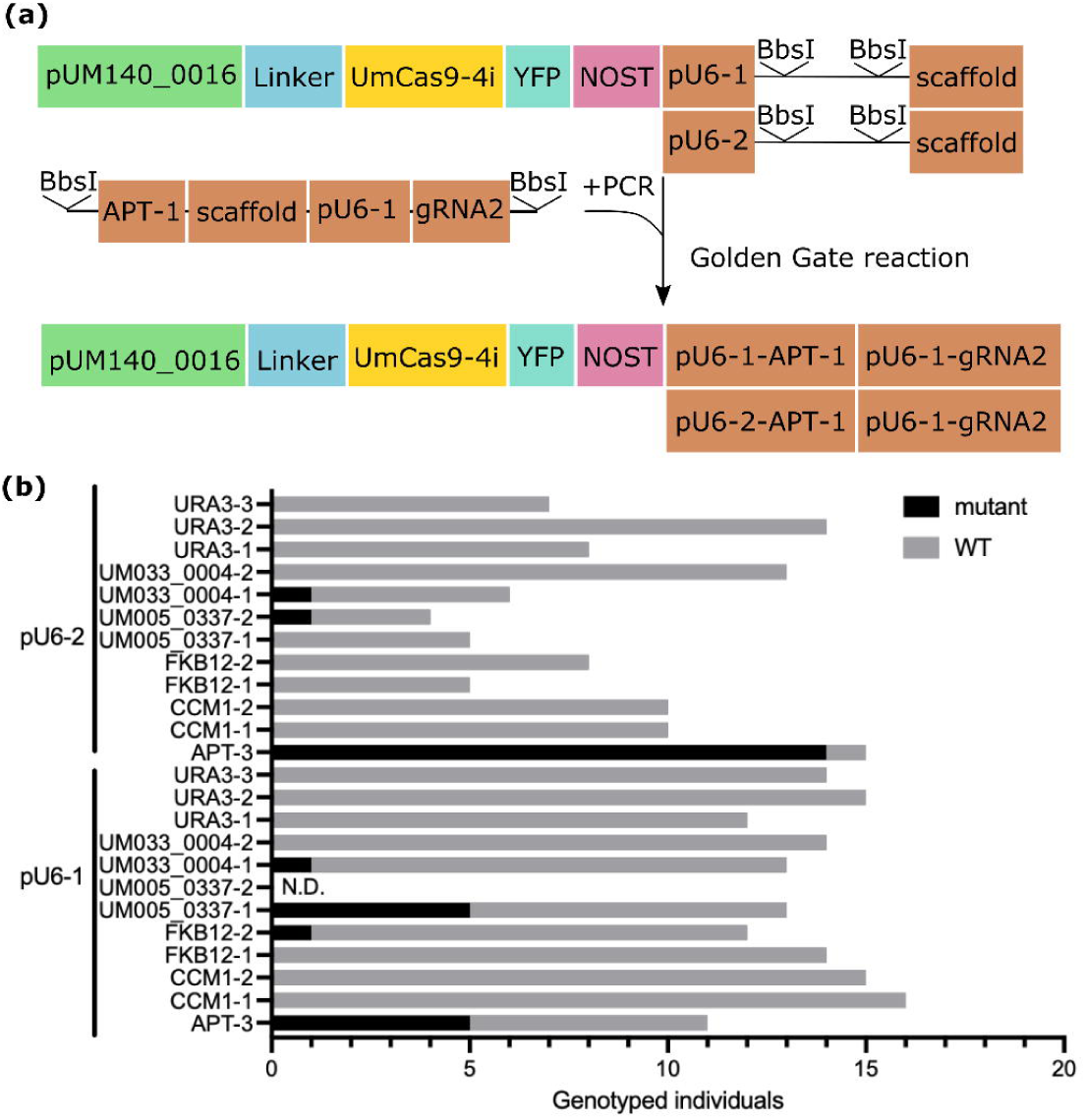
Vector-based co-editing of non-marker genes in *Ulva*. (a) Overview of generating a vector construct targeting APT-1 and a second gene of interest. A PCR-mediated template containing two BbsI recognition sites flanking APT-1-scaffold and pU6-1-gRNA2 is inserted in the destination vector pUM140_0016-UmCas9-4i-YFP-pU6-1-BbsI-scaffold or pUM140_0016-UmCas9-4i- YFP-pU6-2-BbsI-scaffold via a Golden Gate cloning reaction. (b) Summary of genotyping by Sanger sequencing. Four to 16 2-FA resistant individuals were genotyped for in total 12 target sites (APT-3, CCM1-1, CCM1-2, FKB12-1, FKB12-2, UM005_0337-1, UM005_0337-2, UM033_0004-1, UM033_0004-2, URA3-1, URA3-2 or URA3-3), with the gRNA targeting APT-1 under control of either pU6-1 or pU6-2. Number of genotyped individuals is visualized per line, including the mutant and non-edited (WT) individuals. N.D. = not determined.

### Multiplex vector-based Cas9 editing

The vector-based Cas9 experiments demonstrated that the co-editing approach could work and that at least four other targets are functional in addition to APT: FKB12-2, UM005_0337-1, UM005_0337-2 and UM033_0004-1. We next tested if we can mutate more than two genes simultaneously. We made new destination vectors containing Cas9-4i or Cas9-10i directly fused to YFP or not and a module that allows the cloning of six entry vectors (A-ccdB/cat-G) using a single Golden Gate reaction (**Fig. S13a**; (Decaestecker *et al*., 2019)). We selected APT-1 and three other functional targets (FKB12-2, UM005_0337-1 and UM033_0004-1), transformed the vectors in *Ulva* and selected on 2-FA. In total, 47 resistant individuals were recovered (2-17 per construct) and the target sites were genotyped by Sanger sequencing. 23 individuals had a single mutation in APT-1. Only one individual contained mutations in two genes, APT-1 and UM005_0337-1 (**Fig. S13b**). Curiously, two individuals with a mutation in UM033_004-1 or UM005_0337-1 were identified where the APT-1 target was not edited. Together, these results show that additional optimizations are necessary for multiplexing.

### Simplex vector-based Cas12a editing

Both Cas9 and LbCas12a are functional nucleases in RNP-mediated genome editing of *Ulva*. In line with the vector-based Cas9 experiments, we tested if APT can be edited by LbCas12a using vector delivery. First, the promoter region of an *Ulva* U3 gene (UM038_0054) was cloned in the pGG-F entry module including a truncated tRNA fused to Mature Direct Repeat together with a ccdB-cat cassette flanked by AarI restriction enzyme sites. This allows for direct cloning of crRNAs flanked by a Mature Direct Repeat and a PolyT sequence in a one-step cloning reaction using two oligos (**Fig. S14a**; (Gaillochet *et al*., 2023)). Expression vectors were assembled with one of the two gRNAs that were successful in RNP experiments and three nuclease variants: LbCas12a, a version codon optimized for wheat (TaLbCas12a) or *Ulva*, the latter containing 9 RbcsI sequences (UmLbCas12a-9i). We cloned these nucleases with and without direct YFP fusion. 0-11 individuals were resistant to 2-FA, depending on the construct (mutation frequency 8 E^-08^ – 9 E^-07^). No clear edits at the target sites were observed with UmLbCas12a9i or LbCas12a, however indels were observed outside the target site in one line of each variant. Target-specific edits were observed with the TaLbCas12a nuclease (**Fig. S14b**). Similar to the RNP experiments, APT-1 appears to be more efficient compared to APT-2 in the TaLbCas12a vector configurations. The main editing outcomes were deletions at the target site of 3-18 bp, but six out of 20 genotyped individuals were escapes with no edits at the target site. These results show that vector-mediated genome editing using TaLbCas12a is possible, but the efficiency is low compared to RNPs and further optimization is necessary.

## Discussion

*Ulva mutabilis/compressa* is developing into a functional biology model. *Ulva* species are notoriously difficult to distinguish morphologically and we have no certainty of the actual identity of the species used in many historical biological studies (Tran *et al*., 2022). A notable exception is the lab strain, isolated as “*Ulva mutabilis*” in the 1950s, but appeared to be conspecific with *Ulva compressa* sixty years later (Føyn, 1958; Steinhagen *et al*., 2018). Thanks to the continuous cultivation under laboratory conditions, we have a good historical record of studies on *Ulva* development, reproduction and biology (reviewed in (Wichard *et al*., 2015; Blomme *et al*., 2023; Wichard, 2023)). More recently, these studies have been supplemented with a reference genome (De Clerck *et al*., 2018), transcriptome (Liu *et al*., 2022) and transgenic tools (Oertel *et al*., 2015; Blomme *et al*., 2021). Here, we report two important additional steps: the development of novel selection markers and CRISPR/Cas-based targeted genome editing.

Our chemical screen largely confirmed that *Ulva* is tolerant to many different types of selection with a variety of modes-of-action including inhibition of DNA, amino acid or carotenoid synthesis or translation (Oertel *et al*., 2015). We demonstrated that 2-FA and Blasticidin and their corresponding susceptibility and resistance genes, respectively, are functional in *Ulva*. Sulfadiazine, norflurazon and/or SMM could be useful for the generation of future additional selectable markers. Blasticidin is a commonly used antibiotic for selection of mammalian cells (Izumi *et al*., 1991; Kimura *et al*., 1994) and it is also used for various microalgae (Buck *et al*., 2018; Ortega-Escalante *et al*., 2019; de Carpentier *et al*., 2020; Poliner *et al*., 2020; Yin & Hu, 2021; Fujiwara *et al*., 2021; Sprecher *et al*., 2023).

Given the distinct mechanism of Blasticidin (translation inhibition) compared to Phleomycin (generating DNA double-stranded breaks), both antibiotics can be used independently from each other and can be stacked. No cross-reactivity has been observed between Blasticidin and Zeocin, a member of the bleomycin/phleomycin family of chemicals (Buck *et al*., 2018; de Carpentier *et al*., 2020). Therefore, this antibiotic resistance marker could be utilized to supertransform an existing transgenic *Ulva* line with a second construct. One possible application for this strategy is the development of CRISPR assays where cells are transformed with a construct containing both a functional resistance gene and a disrupted second resistance marker. After selection using the first marker, the resistant cells are supertransformed with CRISPR components and a repair template to restore functionality of the second marker, as demonstrated in *Chlamydomonas* (Greiner *et al*., 2017). The cost of Blasticidin is about twice that of Phleomycin, which is a drawback of this selection method.

CRISPR is currently the most effective way to generate targeted DNA mutations. For *Ulva,* it is a more direct way to study gene function in contrast to random mutagenesis (*e.g.* UV radiation and DNA insertion; (Fjeld, 1971; Løvlie, 1978; Oertel *et al*., 2015; Kwantes & Wichard, 2022)). Despite considerable advancements in this technology in many land plant model systems, CRISPR has been reported in only a handful of algae. In seaweeds in particular, Cas9 RNP-based mutagenesis of the APT marker gene is reported in three species: *Ectocarpus sp.7*, *Saccharina japonica* and *Ulva prolifera* (Badis *et al*., 2021; Ichihara *et al*., 2021; Shen *et al*., 2023). Here, we confirm that APT mutagenesis and 2-FA selection is an effective way of evaluating CRISPR efficiency in *Ulva* using Cas9 and Cas12a RNPs and vectors and we have shown that we can genetically complement mutant lines.

The use of a co-editing strategy to couple APT mutagenesis to CRISPR editing of a second gene of interest had variable success. In the end, besides APT, we obtained mutations in just four genes (CCM1, FKB12, UM005_0337 and UM033_0004) and removed five protein coding genes within a 20 kb genomic DNA region. In *Ectocarpus* three genes were also successfully co-edited with APT (Badis *et al*., 2021), suggesting that this strategy is currently the most efficient to identify seaweed mutants with no known or obvious phenotype. Besides co-editing, we demonstrate that targeted insertion of the BleR cassette is a viable alternative to generate *Ulva* mutants. Although the tools presented in this manuscript allow CRISPR editing for single/double mutant studies, eventually additional optimization to generate higher-order mutants will be needed.

The Ulvophytes are an evolutionarily interesting group of organisms because they display a wide variety of thallus and cellular organizations (Del Cortona *et al*., 2020; Hou *et al*., 2022). *Ulva* is ecologically relevant as a primary producer and for causing nuisance blooms in disturbed ecosystems (Poole & Raven, 1997), but it is also cultivated in (integrated) aquaculture (Bolton *et al*., 2016). Emerging -omics data and molecular tools now allow the study of the effects of gene overexpression or mutation on *Ulva* biology linked to evolution, ecology and industrial applications. The haploid *Ulva* reference genome is relatively small (98,5 Mbp) and contains 12,924 protein coding genes (De Clerck *et al*., 2018). Moving forward, it is clear that an increase in vector-based CRISPR efficiency will be necessary to perform larger-scale experiments. Boosting efficiency should avoid the necessity of co-targeting of APT (Wang *et al*., 2024). We attempted to increase efficiency via a variety of vector configurations. We were able to clearly demonstrate that the U6-1 promoter outperformed U6-2. Since Cas9 is a relatively large transgene (4,1 kb), we hypothesize that constitutive expression is prone to silencing as observed in *Chlamydomonas* (Jiang *et al*., 2014; Neupert *et al*., 2020). In our experiments, codon optimization and including endogenous regulatory elements such as *Ulva* promoters and introns have improved, but not drastically affected, CRISPR efficiency. The frequent insertion of vector DNA in mutant individuals and our targeted insertion experiment suggests that homology-directed repair can become a powerful tool in *Ulva* genome engineering. Similar observations have been made in *Ulva prolifera* and *Chlamydomonas* (Greiner *et al*., 2017; Picariello *et al*., 2020; Akella *et al*., 2021; Ichihara *et al*., 2024) and need to be further explored in our system. Besides vector optimization, other factors such as cultivation conditions can affect mutation frequency (Freudenberg *et al*., 2022) and these should be evaluated in the future.

## Acknowledgements

J.B., T.B.J., and O.D.C. are indebted to Research Foundation—Flanders (FWO) for funding (Senior Research Project G015623N, postdoctoral fellowship 12T3418N, ERC StG runner- project G0AHR24N and ESFRI grants GOH3817N and I001621N). T.B.J. and J.B. thank Ghent University Special Research fund (postdoctoral fellowship BOF20/PDO/016 and starting grant BOF/STA/202209/016) and VIB Tech Watch Project De-Risking Funding. J.B. and O.D.C. thank the Gordon and Betty Moore Foundation (Symbiosis in Aquatic Systems Initiative #9329) The authors thank Thomas Wichard for providing the bacterial strains *Cythopaga sp.* MS6 and *Roseobacter sp.* MS2.

## Author contributions

J.B., O.D.C. and T.B.J. conceived and designed the study. J.B. and J.A.R. generated, analysed and interpreted data. J.B. and T.B.J. led the writing of the manuscript with support of all authors.

## Data availability

The authors declare that the data supporting the findings of this study are available within the paper and its Supporting Information.

## Supporting Information

The Supporting Information consist of 4 tables and 14 figures:

Figure S1. *Ulva* chemical sensitivity assay.

Figure S2. Cytophaga sp. MS6 and Roseobacter sp. MS2 sensitivity assay.

Figure S3. Architecture of destination vectors with integrated Blasticidin resistance cassette.

Figure S4. Schematic representation of target genes.

Figure S5. Codon optimized Cas9 is functional in the model plant Arabidopsis.

Figure S6. *In vitro* Cas9 digest.

Figure S7. RNP editing of the Wild-type strain.

Figure S8. RNP dilution experiment.

Figure S9. *Ulva* targeted genome editing with Cas12a RNPs.

Figure S10. Genetic complementation of *Ulva apt* mutants.

Figure S11. Targeted insertion of BleR in CCM1 and URA3.

Figure S12. Co-editing with RNPs in *Ulva*.

Figure S13. Vector-based multiplex genome editing in *Ulva*.

Figure S14. Vector based genome editing using Cas12a in *Ulva*.

Table S1. Relative transformation and/or editing efficiency per experiment.

Table S2. List of gRNA and primer sequences used in this study

Table S3. Complete list of generated vectors (entry, destination and expression).

Table S4. Summary of large deletion experiment.

