## Supplementary material for "Consolidating *Ulva* functional genomics: Gene editing and new selection systems": Ulva_CRISPR_SI_v6.pdf

The Supporting Information consist of 4 tables and 14 figures:

**Figure S1.** *Ulva* chemical sensitivity assay.
**Figure S2.** *Cytophaga sp.* MS6 and *Roseobacter sp.* MS2 sensitivity assay.
**Figure S3.** Architecture of destination vectors with integrated Blasticidin resistance cassette.
**Figure S4.** Schematic representation of target genes.
**Figure S5.** Codon optimized Cas9 is functional in the model plant Arabidopsis.
**Figure S6.** *In vitro* Cas9 digest.
**Figure S7.** RNP editing of the Wild-type strain.
**Figure S8**. RNP dilution experiment.
**Figure S9.** *Ulva* targeted genome editing with Cas12a RNPs.
**Figure S10.** Genetic complementation of *Ulva apt* mutants.
**Figure S11.** Targeted insertion of BleR in CCM1 and URA3.
**Figure S12.** Co-editing with RNPs in *Ulva*.
**Figure S13.** Vector-based multiplex genome editing in *Ulva*.
**Figure S14.** Vector based genome editing using Cas12a in *Ulva*.

**Table S1.** Relative transformation and/or editing efficiency per experiment.
**Table S2.** List of gRNA and primer sequences used in this study
**Table S3.** Complete list of generated vectors (entry, destination and expression).
**Table S4.** Summary of large deletion experiment.

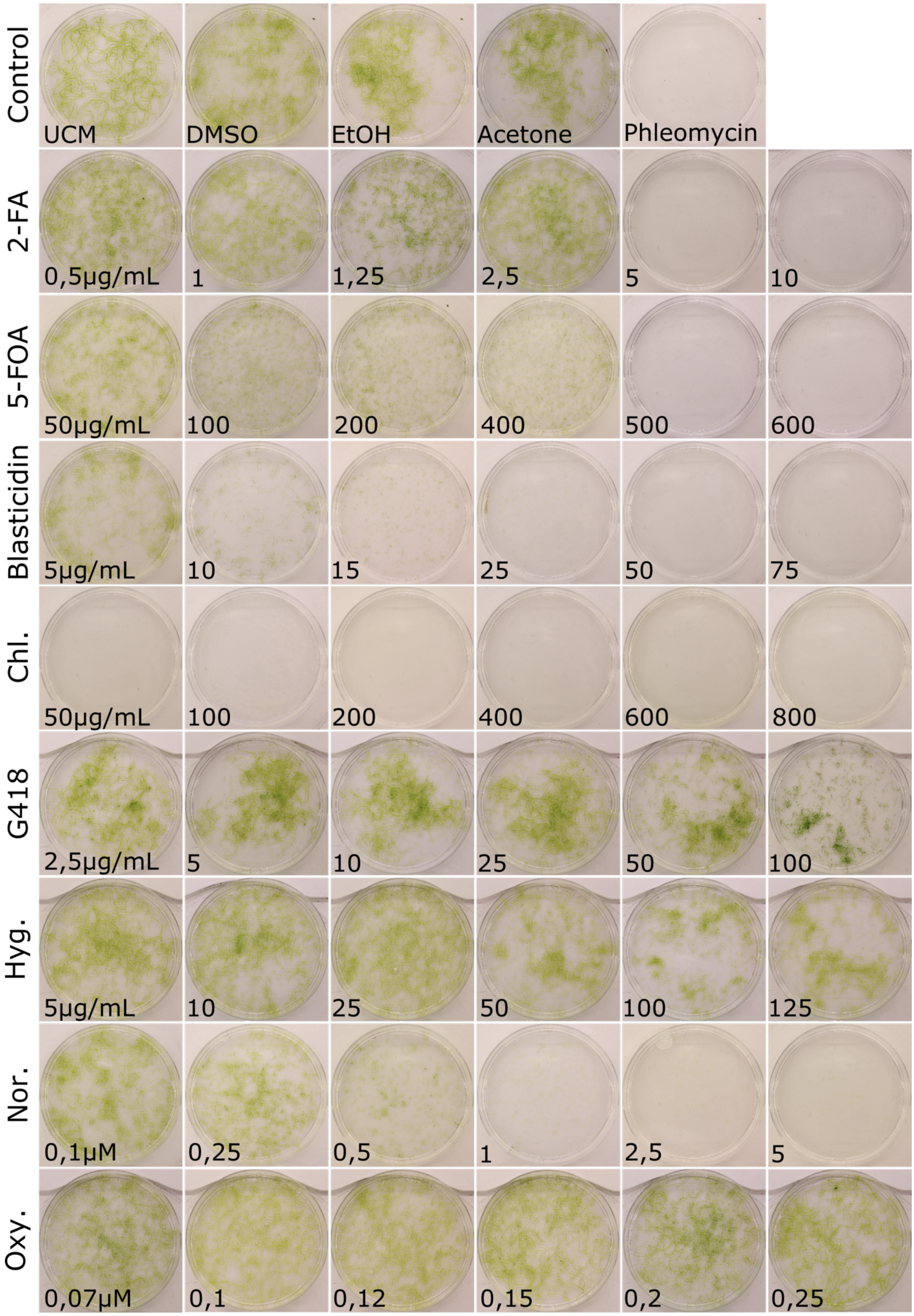
Control
UCM
DMSO
EtOH
Acetone
Phleomycin
2-FA
0,5µg/mL
1
1,25
2,5
5
10
5-FOA
50µg/mL
100
200
400
500
600
Blasticidin
5µg/mL
10
15
25
50
75
Chl.
50µg/mL
100
200
400
600
800
G418
2,5µg/mL
5
10
25
50
100
Hyg.
5µg/mL
10
25
50
100
125
Nor.
0,1µM
0,25
0,5
1
2,5
5
Oxy.
0,07µM
0,1
0,12
0,15
0,2
0,25

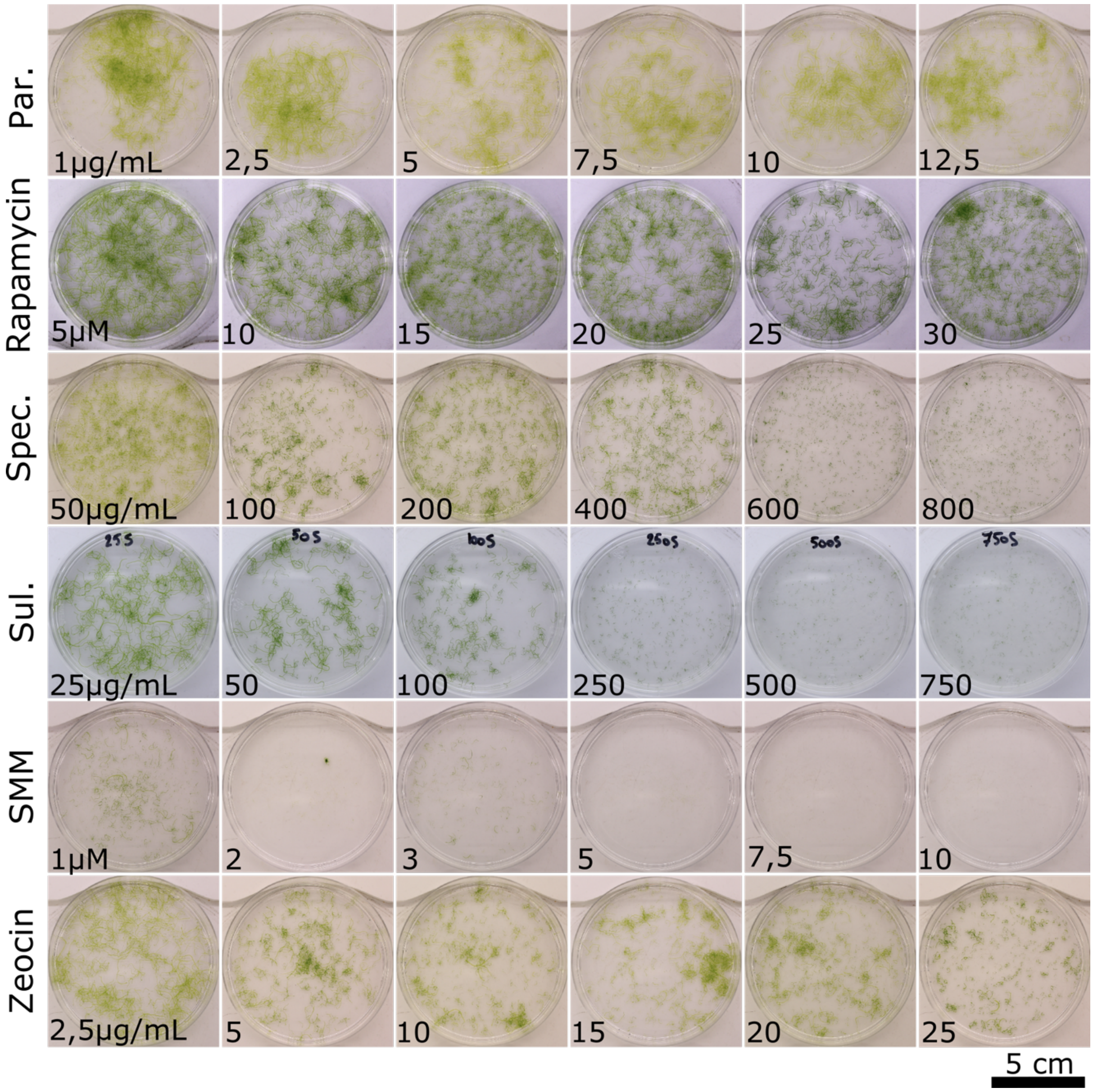


**Figure S1.** ***Ulva*** **chemical sensitivity assay.**
*Ulva* germlings were transferred to increasing concentrations of 2-Fluoroadenine (2-FA), 5-Fluoro-orotic acid (5-FOA), Blasticidin, Chloramphenicol (Chl.), G418, Hygromycin (Hyg.), Norflurazon (Nor.), Oxyfluorfen (Oxy.), Paromomycin (Par.), Rapamycin, Spectinomycin (Spec.), Sulfadiazin (Sul.), Sulfometuron Methyl (SMM) and Zeocin. Cultures were photographed two weeks after transfer. Controls are solvents used to dissolve the chemicals and 50 µg/mL Pheomycin. All values represent concentrations in µg/mL or µM. Scale = 5cm.

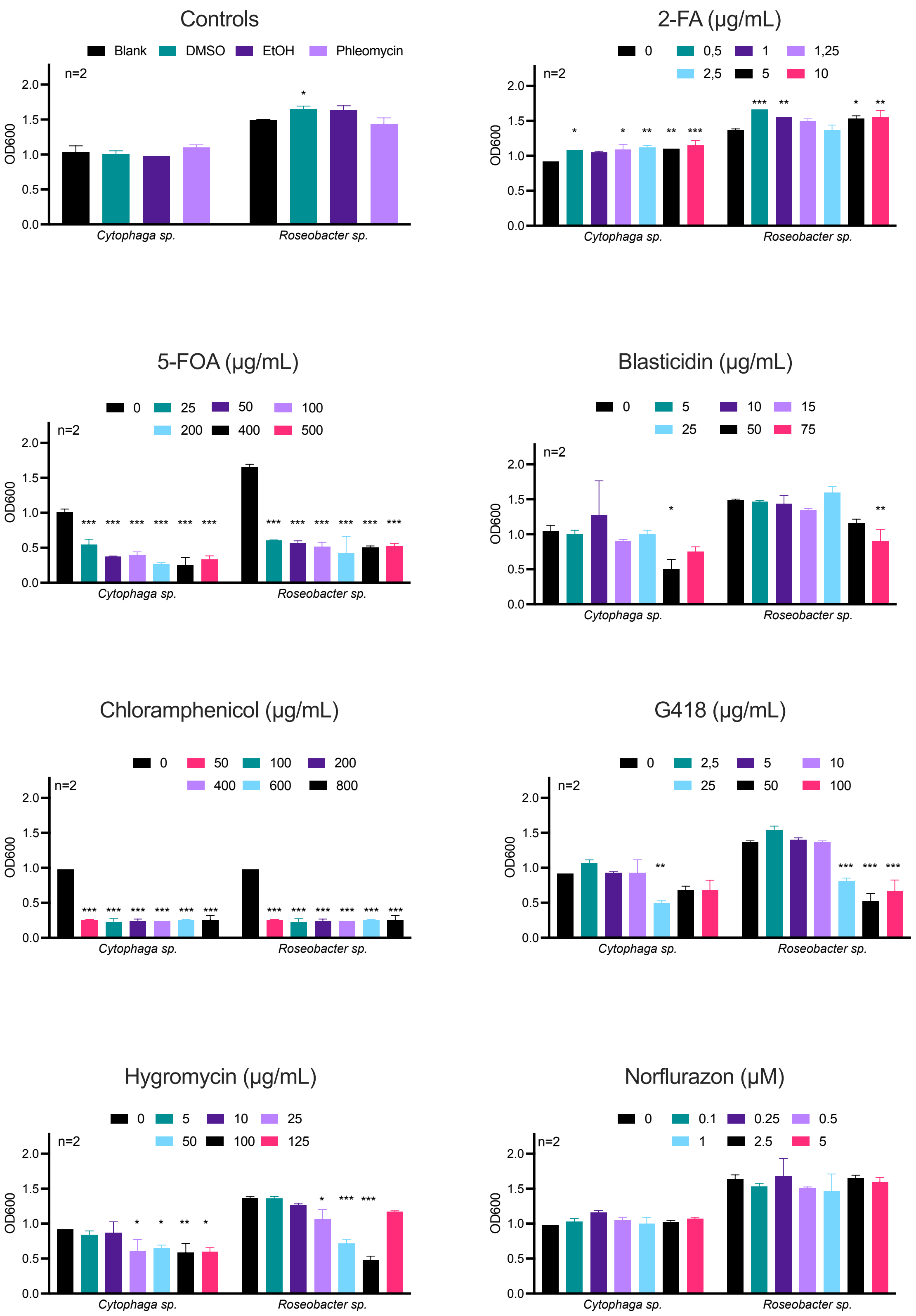

Controls
Blank
DMSO
EtOH
Phleomycin
n=2
OD600
2.0
1.5
1.0
0.5
0.0
*Cytophaga sp.*
*Roseobacter sp.*
2-FA (µg/mL)
0
0,5
1
1,25
2,5
5
10
n=2
OD600
*Cytophaga sp.*
*Roseobacter sp.*
5-FOA (µg/mL)
0
25
50
100
200
400
500
n=2
OD600
*Cytophaga sp.*
*Roseobacter sp.*
Blasticidin (µg/mL)
0
5
10
15
25
50
75
n=2
OD600
*Cytophaga sp.*
*Roseobacter sp.*
Chloramphenicol (µg/mL)
0
50
100
200
400
600
800
n=2
OD600
*Cytophaga sp.*
*Roseobacter sp.*
G418 (µg/mL)
0
2,5
5
10
25
50
100
n=2
OD600
*Cytophaga sp.*
*Roseobacter sp.*
Hygromycin (µg/mL)
0
5
10
25
50
100
125
n=2
OD600
*Cytophaga sp.*
*Roseobacter sp.*
Norflurazon (µM)
0
0.1
0.25
0.5
1
2.5
5
n=2
OD600
*Cytophaga sp.*
*Roseobacter sp.*

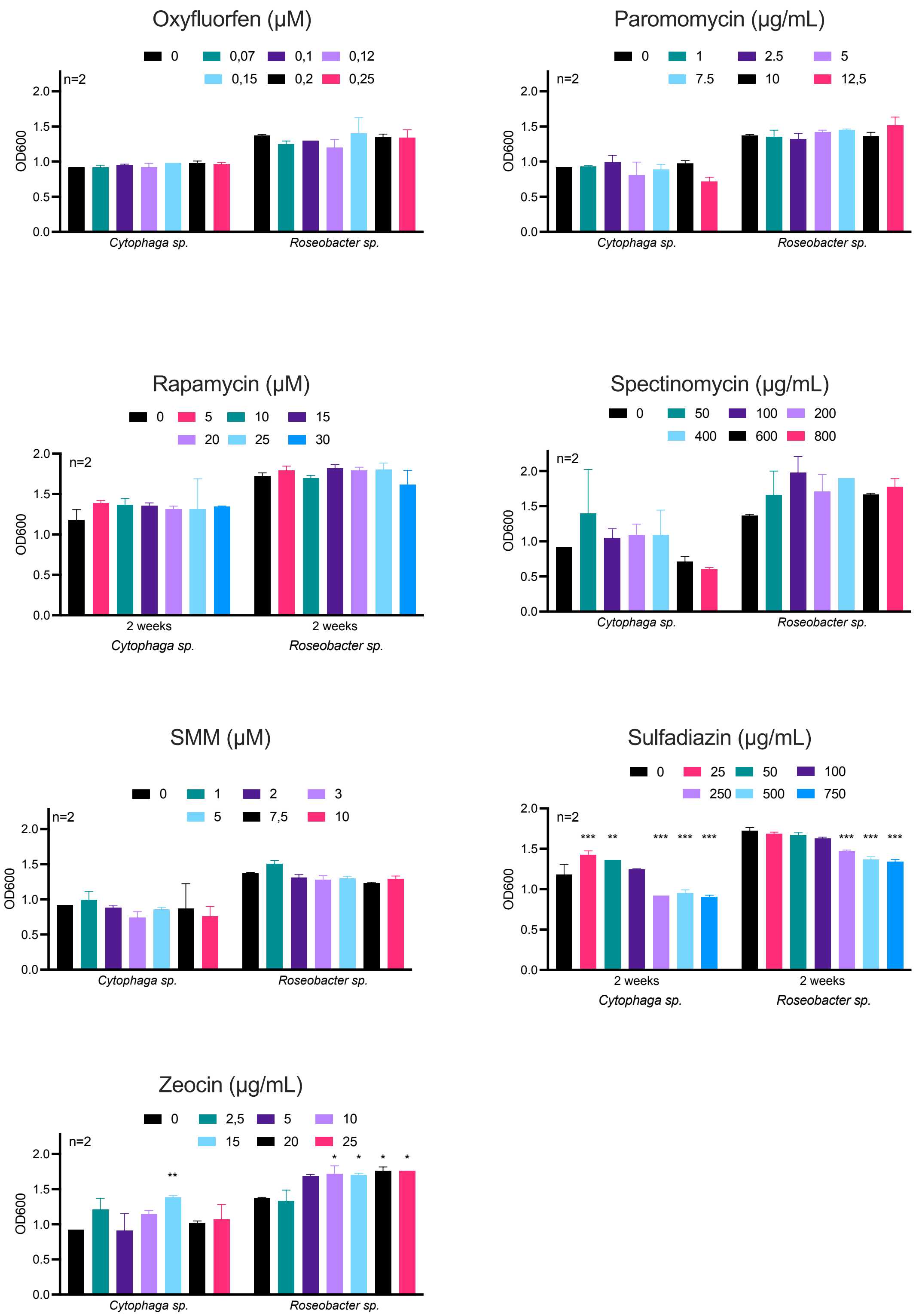


**Figure S2. *Cytophaga sp.* MS6 and *Roseobacter sp.* MS2 sensitivity assay.**
*Cythopaga sp.* MS6 and *Roseobacter sp.* MS2 were inoculated at increasing concentrations of 2-Fluoroadenine (2-FA), 5-Fluoro-orotic acid (5-FOA), Blasticidin, Chloramphenicol, G418, Hygromycin, Norflurazon, Oxyfluorfen, Paromomycin, Spectinomycin, Sulfometuron Methyl (SMM) and Zeocin. All values represent average OD600 ± SD (n=2). Controls are solvents used to dissolve the chemicals and 50 µg/mL Pheomycin. Significant differences from the blank control are indicated (ANOVA with *post hoc* Dunnett multiple comparison test; * = P >0,05; ** = P >0,01 and *** = P >0,001).

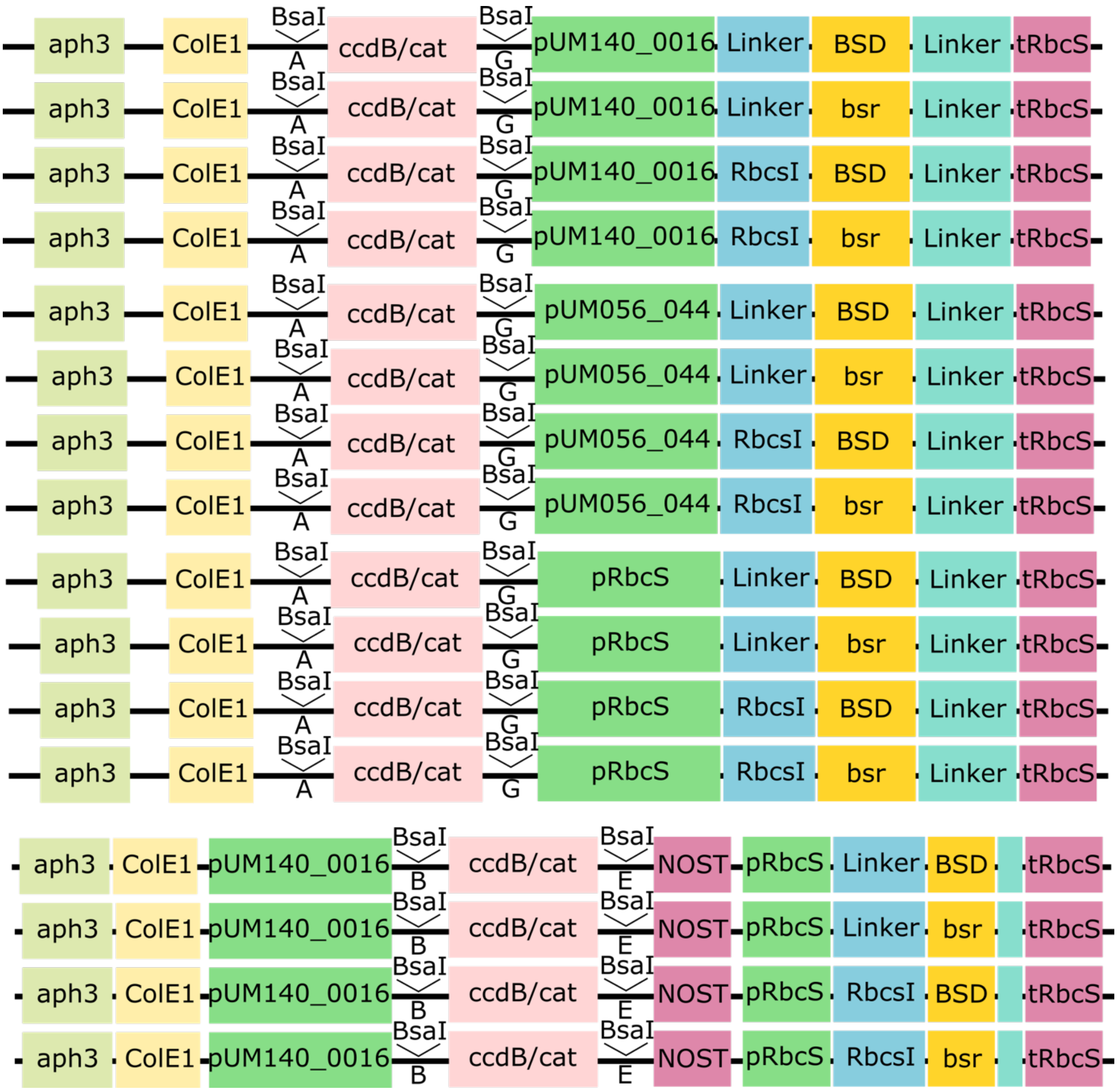


**Figure S3. Architecture of destination vectors with integrated Blasticidin resistance cassette.**

Standard destination vectors contain an ACCT (“A”) and GTAT (“G”) Golden Gate overhang after BsaI digest. BSD or bsr is under control of one of three promoters (pUM140_0016, pUM056_0044 or pRbcS) and a RbcsI sequence or not. All pRbcS constructs driving BSD or bsr were integrated in pUM140_0016-B-ccdB/cat-E-NOST destination vectors containing an AACA (“B”) and CTGC (“E”) overhang that facilitate the generation of tagged lines. All vectors are available on https://vectorvault.vib.be. See also **Table S2**.

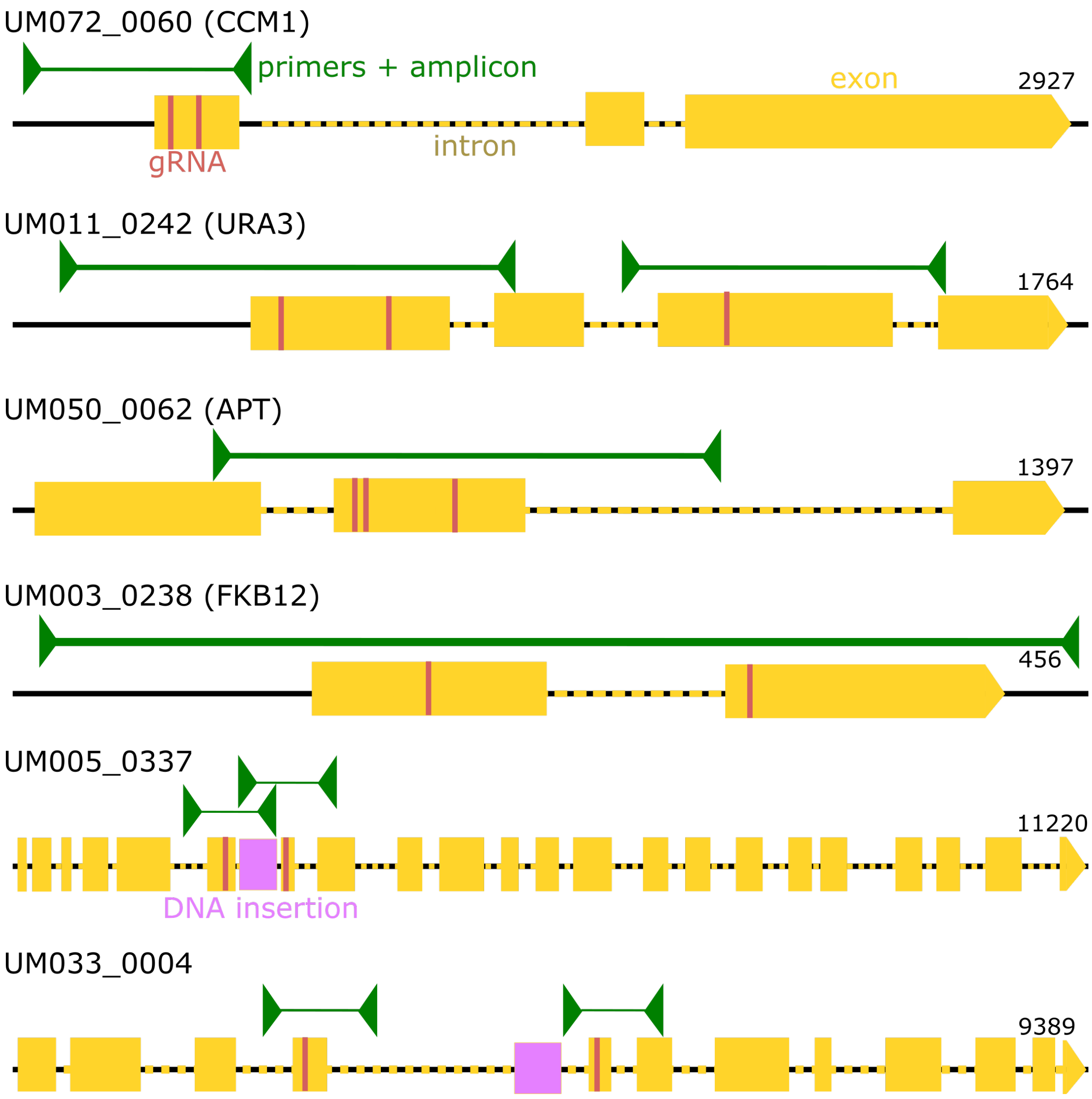


**Figure S4. Schematic representation of target genes.**
Gene models of the target genes with exons in yellow, introns in dashed yellow line, gRNAs in red and genotyping primers with amplicons in green. For UM005_0337 and UM033_0004 the DNA insertion reported in (Kwantes & Wichard, 2022) is annotated in a magenta. Gene length not to scale, the length of gene including introns is indicated. gRNA and primer sequences are provided in **Table S2**.

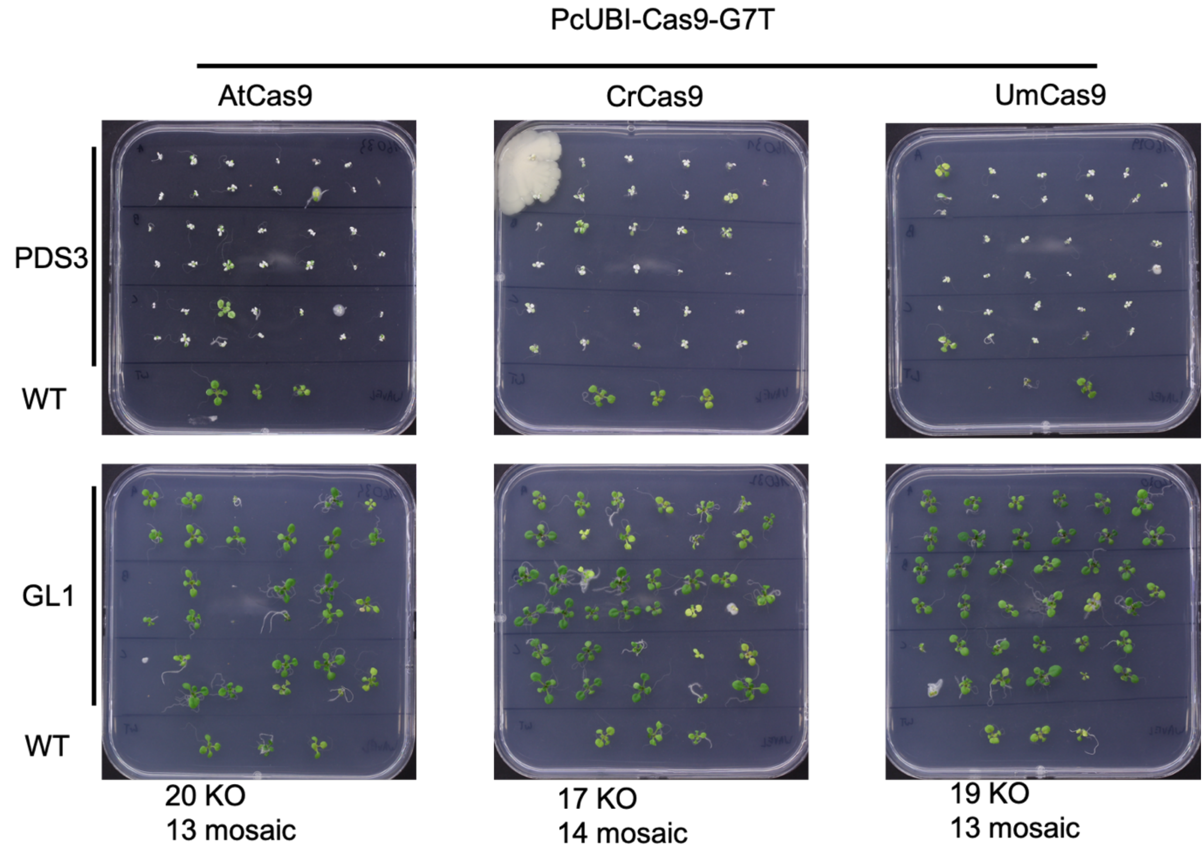


**Figure S5. Codon optimized Cas9 is functional in the model plant Arabidopsis.**
A Cas9 nuclease codon optimized for *Arabidopsis* (AtCas9), *Chlamydomonas* (CrCas9) and *Ulva* (UmCas9) was generated. The variants were transformed in Arabidopsis, targeting either PHYTOENE DESATURASE3 (PDS3) or GLABRA1 (GL1). *pds3* mutants are albino and *gl1* mutants have no trichomes. Similar numbers of T1 plants with mosaic or KO *pds3* or *gl1* phenotypes were observed for UmCas9, CrCas9 and AtCas9. The number of mosaic or KO *glabrous* individuals are noted beneath the respective plates. For each construct a wild-type (WT) control is included.

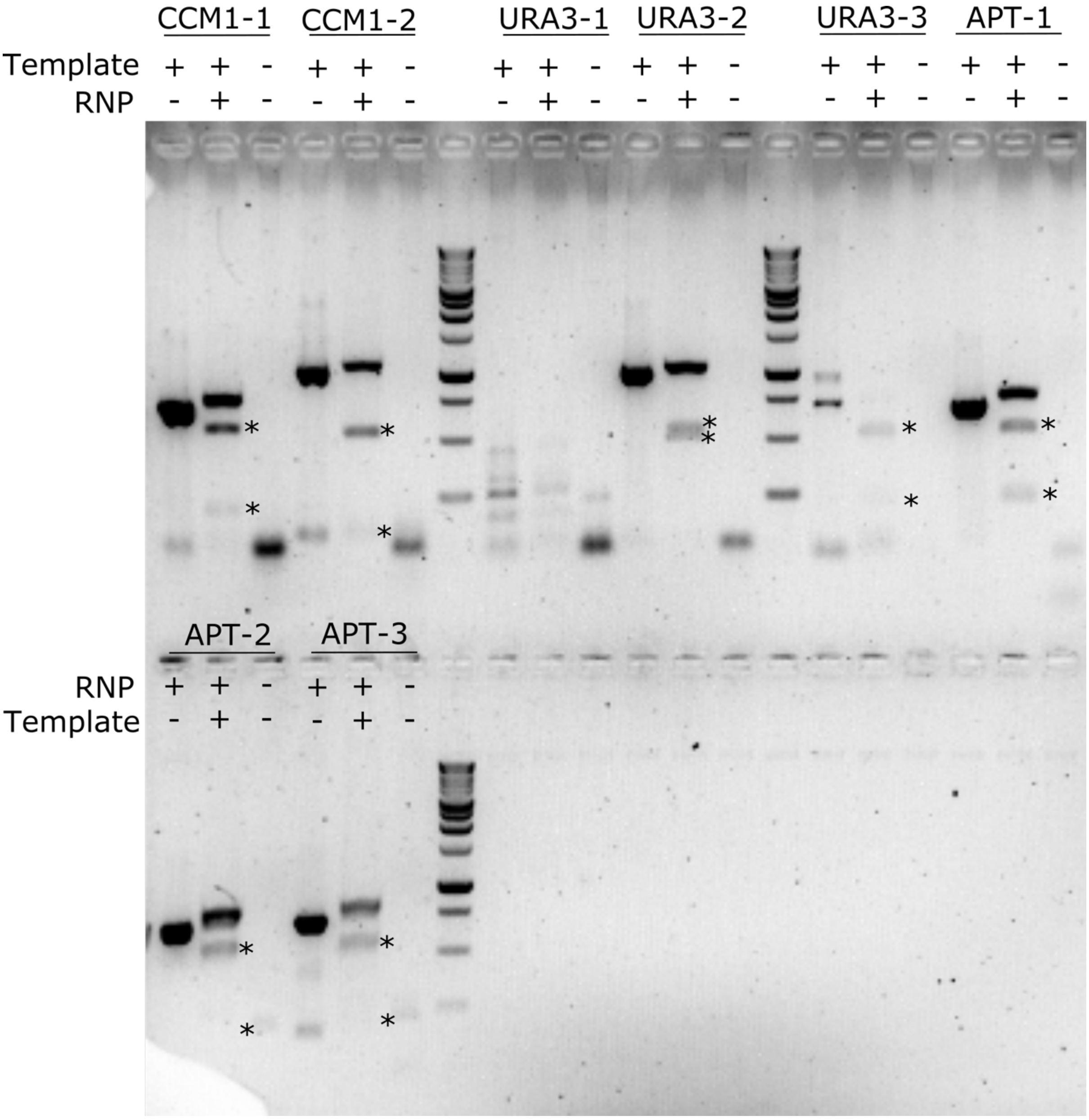


**Figure S6.** ***In vitro*** **Cas9 digest.**
All target sites (CCM1-1, CCM1-2, URA3-1, URA3-2, URA3-3, APT-1, APT-2 and APT-3) were amplified from gDNA. The amplicons were incubated (+) or not (-) with the respective RNP. A no-template water control sample is included. Expected size of the digested amplicon are indicated with an asterisk. DNA marker is Benchtop 1kb ladder (Promega).

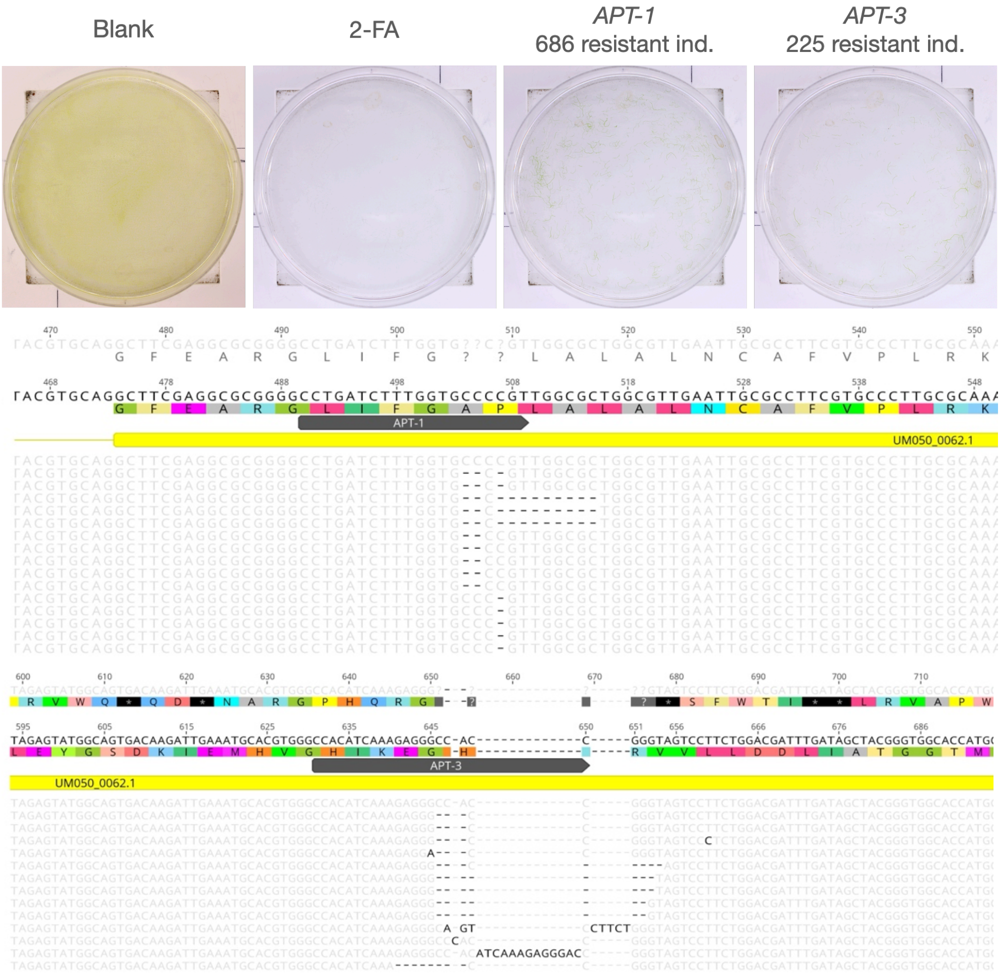


**Figure S7. RNP editing of the Wild-type strain.**
The Wild-type strain can be edited at APT-1 and APT-3 target site. A control plate containing non-selective medium (Blank) and 2-FA control is included. The number of 2-FA resistant individuals is indicated above each transfection plate. Below the figure are observed indels in 2-FA resistant individuals. Sanger sequencing reads were aligned to the reference gene sequence using Geneious software. The target site (APT-1 or APT-3) is annotated in grey on the coding sequence (yellow). Every read represents one genotyped individual. An untransfected control is included (first read).

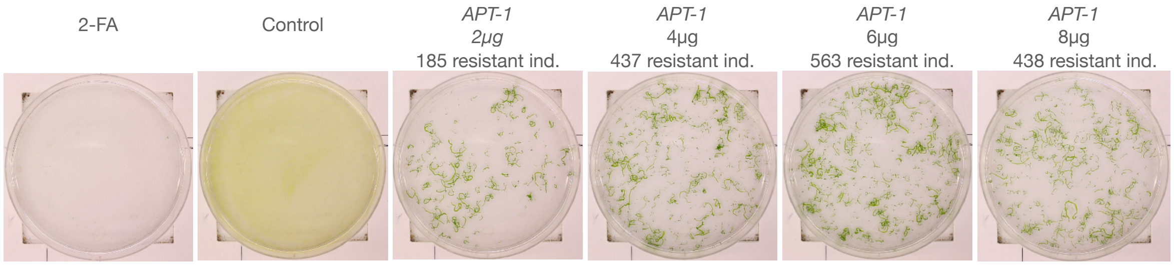


**Figure S8**. **RNP dilution experiment.**
A reduction in RNP concentration still leads to numerous 2-FA resistant *Ulva* individuals. A control plate containing non-selective medium (Blank) and 2-FA control is included. The number of 2-FA resistant individuals is indicated above each transfection plate with the respective APT-1 RNP Cas9 concentration in µg.

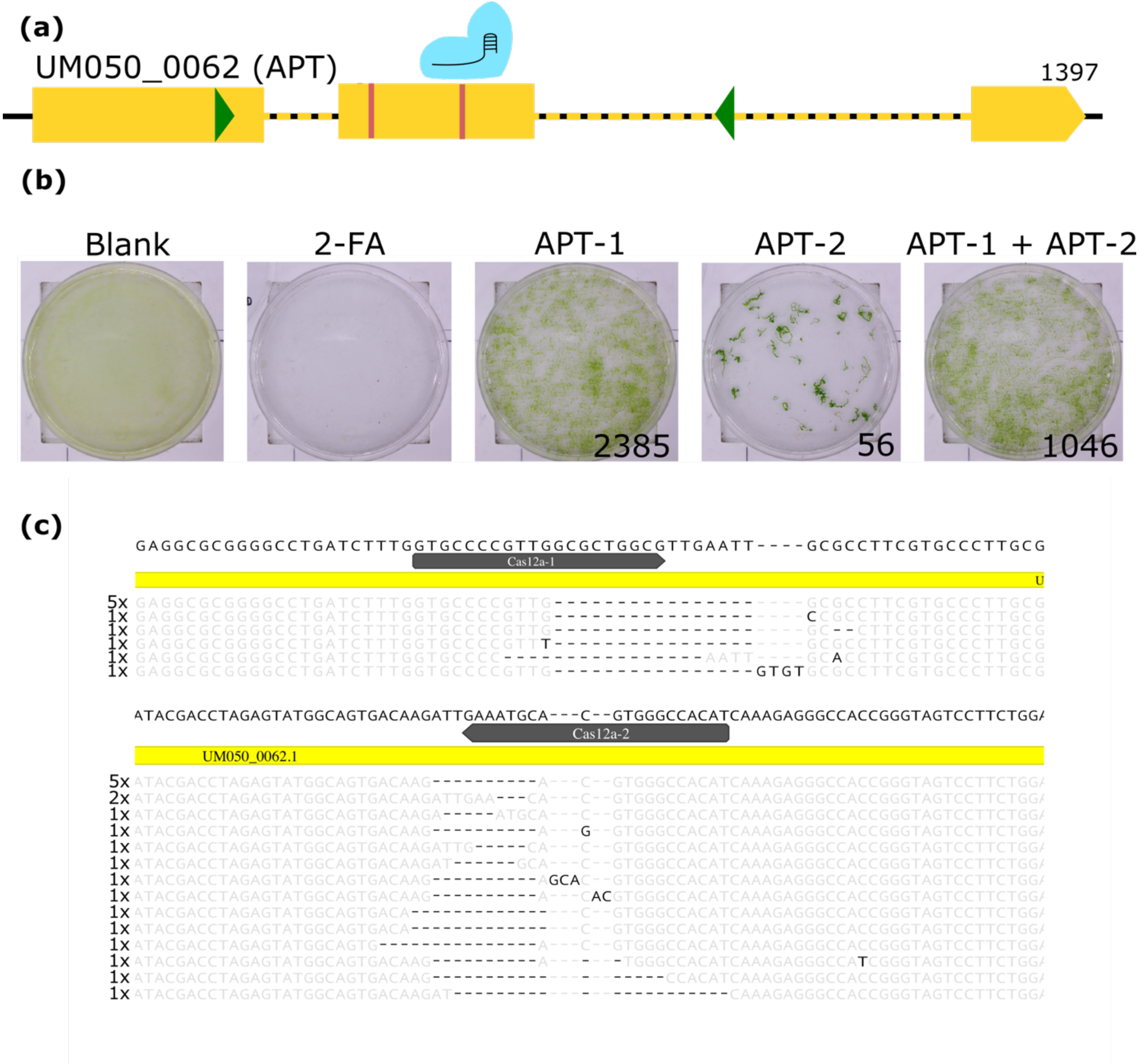


**Figure S9.** ***Ulva*** **targeted genome editing with Cas12a RNPs.**

(a) Overview of experiment. The Cas12a RNP targeting APT-1 and/or APT-2 was transfected and resistant individuals were selected by adding 2-FA. Green arrowheads indicate genotyping primers.

(b) Result of Cas12a RNP transfection and 2-FA selection. Number of resistant individuals is indicated on the transfection plates.

(c) Observed indels in 2-FA resistant individuals. Sanger sequencing reads were aligned to the reference gene sequence. The target site (APT-1 and APT-2) is annotated in grey on the coding sequence (yellow) via Geneious software. Every mutant read contains the number of genotyped individuals with the same indel pattern.

(a)

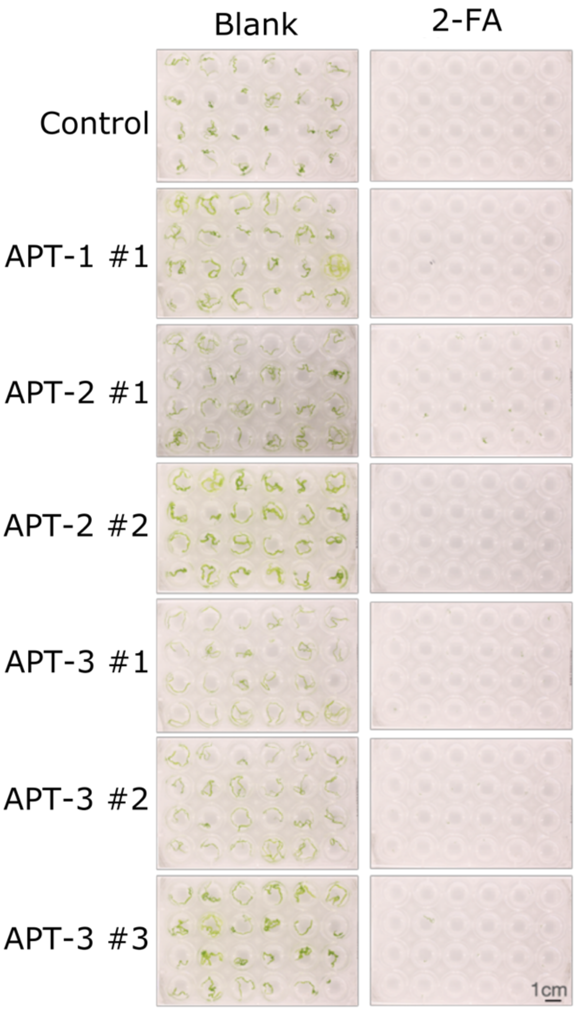


(b)

| Individual | ICE | R Squared |
|---|---|---|
| *APT-1* comp. #1 | 45 | 0,94 |
| *APT-1* comp. #2 | 30 | 0,96 |
| *APT-1* comp. #3 | 29 | 0,97 |
| *APT-1* comp. #4 | 27 | 0,96 |
| *APT-2* comp. #1 | 2 | 0,99 |
| *APT-2* comp. #2 | 2 | 0,99 |
| *APT-2* comp. #3 | 9 | 0,98 |
| *APT-2* comp. #4 | 16 | 0,98 |
| *APT-3* comp. #1 | 46 | 0,96 |
| *APT-3* comp. #2 | 51 | 0,96 |
| *APT-3* comp. #3 | 41 | 0,97 |
| *APT-3* comp. #4 | 47 | 0,96 |
| *APT-3* comp. #5 | 9 | 0,98 |
| *APT-3* comp. #6 | 11 | 0,98 |
| *slender* control #1 | 0 | 1 |
| *slender* control #2 | 0 | 1 |

**Figure S10. Genetic complementation of *Ulva apt* mutants.**

(a) Mutants for three different APT target sites (APT-1, APT-2 and APT-3) were transformed with pUM140_0016-APT-YFP and propagated. The progeny of these transformants was cultured in standard UCM medium or UCM supplemented with 2-FA. Scale: 1cm

(b) Genotyping by Sanger sequencing of complemented lines. Table shows the ICE analysis (https://ice.synthego.com). ICE score indicates the indel %, the R squared value indicates how well the proposed indel distribution fits the Sanger sequence data of the edited sample.

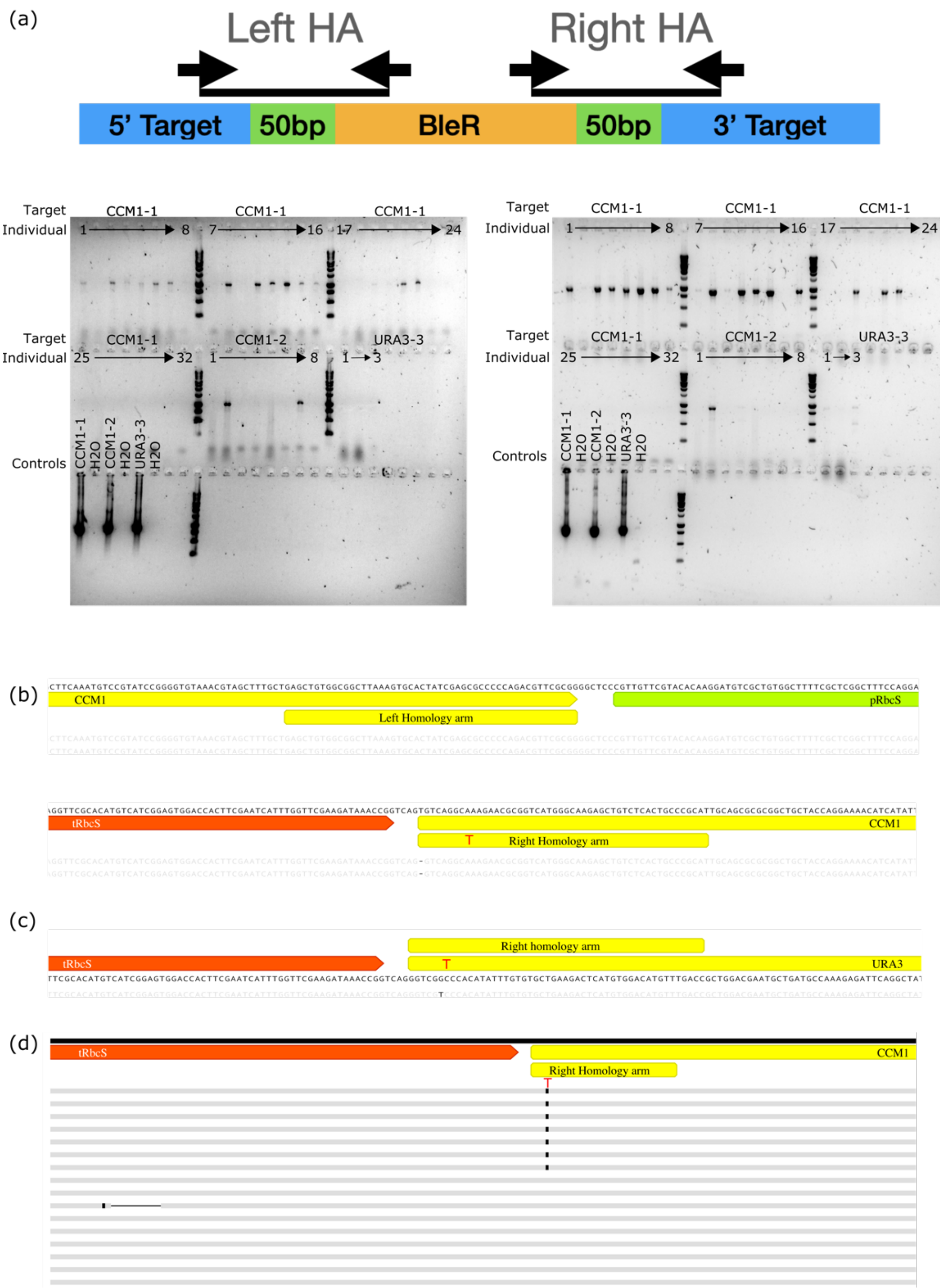


**Figure S11. Targeted insertion of BleR in CCM1 and URA3.**

(a) PCR strategy to amplify the left or right homology arm (HA) is shown above the gel electrophoresis result. Target and individuals are labeled, and the controls are amplification using a plasmid control construct or water ($H_2O$) control. DNA marker is Benchtop 1kb ladder (Promega).

(b) Sanger sequencing result of left and right homology arms for CCM1-2 target. SNP included in the right homology arm is indicated in red font. Visualised with Geneious software.

(c) Sanger sequencing result of right homology arm for URA3-3 target. SNP included in the right homology arm is indicated in red font. Visualised with Geneious software.

(d) Sanger sequencing result of right homology arms for CCM1-1 target. SNP included in the right homology arm is indicated in red font. A 17bp deletion in tRbcS in one read is indicated as a black line. Visualised with Geneious software.

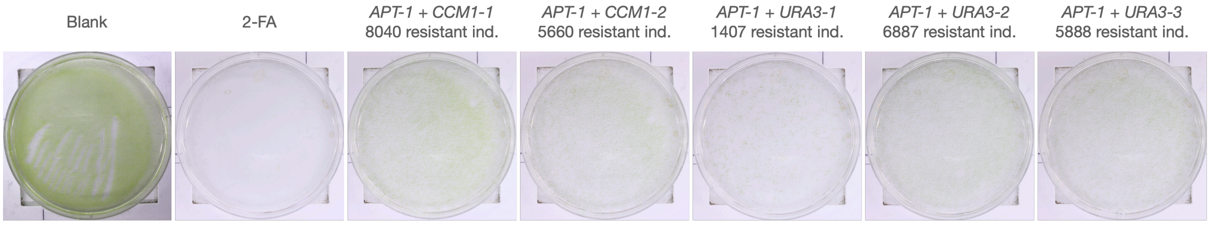


| | APT-1 + CCM1-1 | APT-1 + CCM1-2 | APT-1 + URA3-1 | APT-1 + URA3-2 | APT-1 + URA3-3 |
|---|---|---|---|---|---|
| Individuals Harvested | 96 | 96 | 96 | 96 | 96 |
| Genotyped | 85 | 71 | 73 | 74 | 77 |
| Mutants | 0 | 1 | 0 | 0 | 0 |

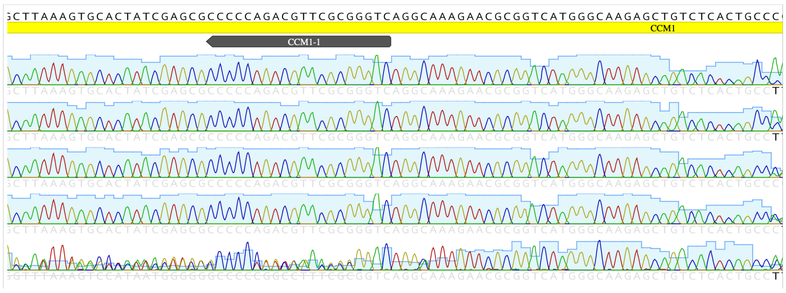


**Figure S12. Co-editing with RNPs in *Ulva*.**
One RNP targeting APT-1 and a second RNP (targeting CCM1-1, CCM1-2, URA3-1, URA3-2 or URA3-3) were co-transfected in *Ulva*. The number of 2-FA resistant individuals are denoted on the transfection plates. The number of harvested and genotyped 2-FA resistant individuals are indicated below each transfection plate, also the number of mutant individuals in the second gene-of-interest is indicated. The Sanger sequencing profile of four non-edited and one mutant individual at the CCM1-1 target site is visualized using Geneious software.

(a)

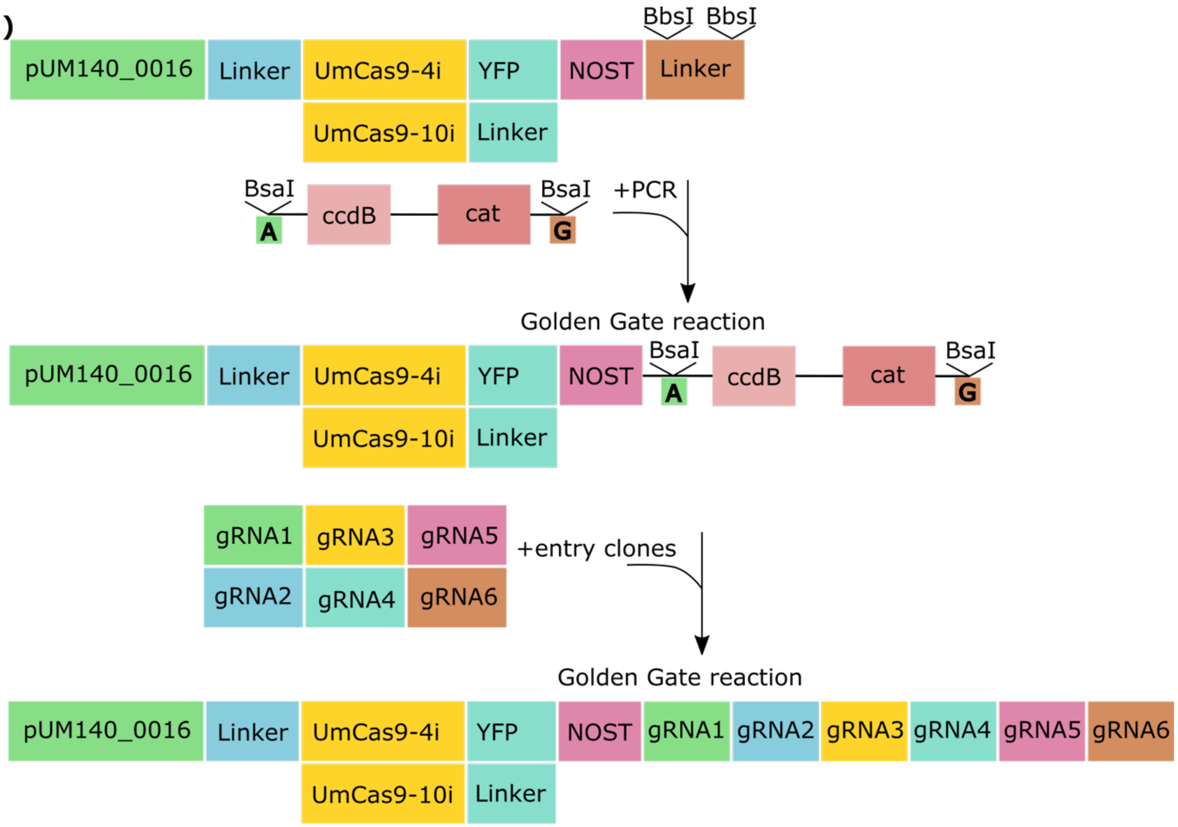


(b)

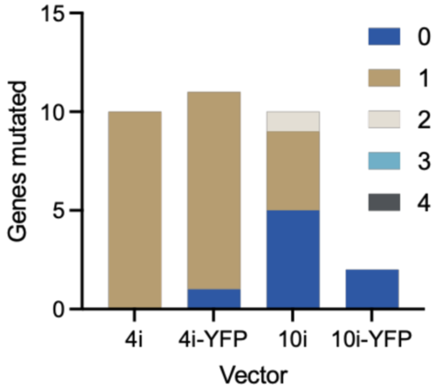


**Figure S13. Vector-based multiplex genome editing in *Ulva*.**

(a) Overview of molecular cloning steps to generate a vector targeting up to six regions of interest. A PCR-mediated template containing two BsaI recognition sites flanking ccdB and chloramphenicol resistance gene (cat) is inserted in the destination vector pUM140_0016-UmCas9-4i-YFP-NOST-BbsI, pUM140_0016-UmCas9-10i-YFP-NOST-BbsI, pUM140_0016-UmCas9-4i-NOST-BbsI and pUM140_0016-UmCas9-10i-NOST-BbsI via a Golden Gate cloning reaction. This new destination vector has an "A" and "G" Golden Gate overhang, allowing to clone up to six entry clones containing a different pU6-gRNA-scaffold construct.

(b) Summary of genotyping by Sanger sequencing. Two to 11 2-FA resistant individuals were genotyped for four different vector combinations: UmCas9-4i or UmCas9-10i fused to YFP or not. All vectors contained APT-1, FKB12-2, UM033_004-1 and UM005_0337-2 target sites. The number of mutated genes per individual is visualized per line.

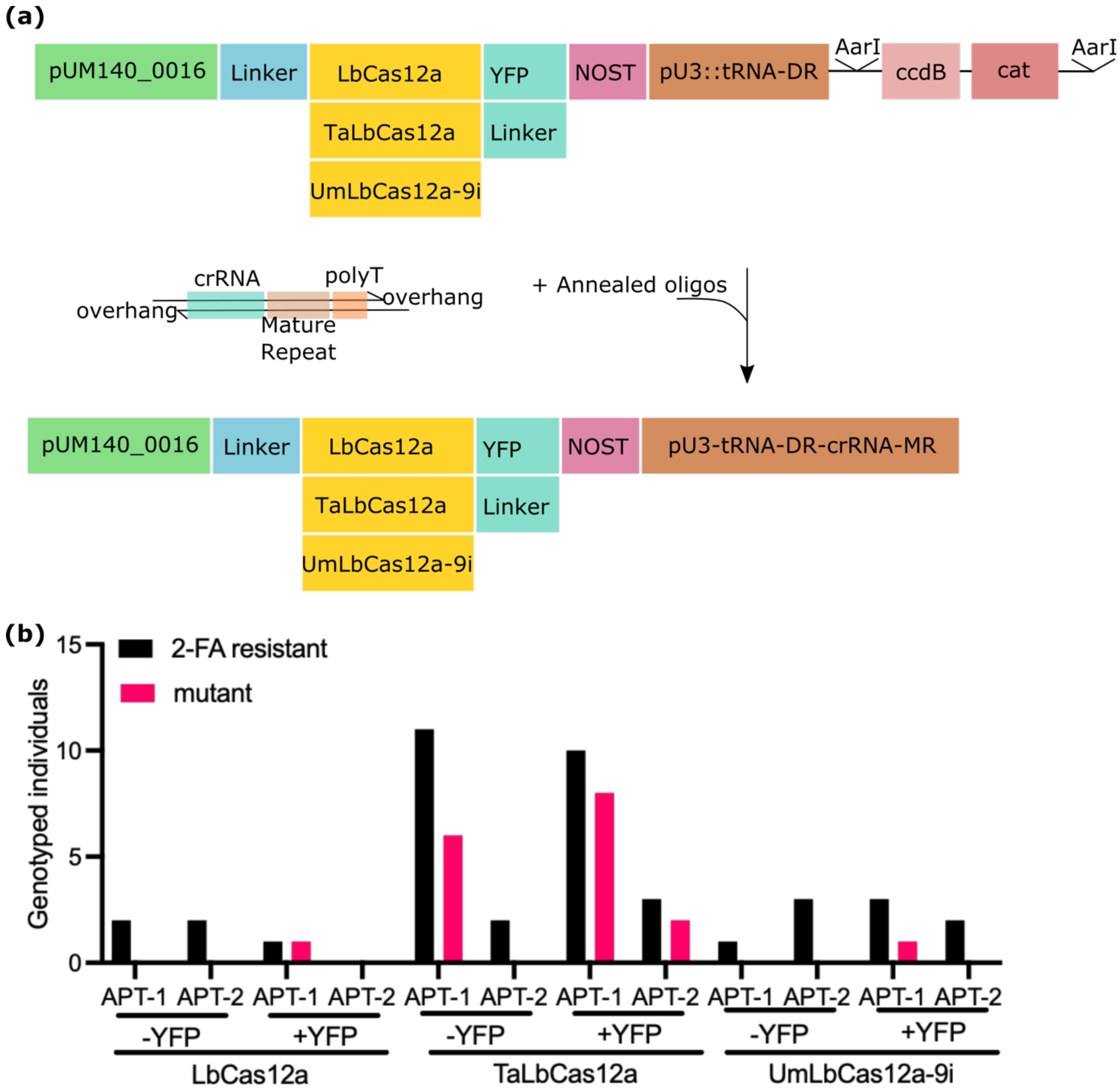


**Figure S14. Vector based genome editing using Cas12a in *Ulva*.**

(a) Overview of generating a vector construct containing one of three different codon optimized LbCas12a sequences: Arabidopsis (LbCas12a), wheat (TaLbCas12a) or *Ulva* containing 9 RbscI sequences (UmLbCas12a-9i). An *Ulva* U3 promoter was cloned including a truncated tRNA fused to Mature Direct Repeat together with a ccdB/cat cassette flanked by AarI restriction enzyme sites. This allows to directly clone in crRNAs flanked by a Mature Direct Repeat and a PolyT sequence in a one-step cloning reaction using two oligos (Decaestecker *et al.*, 2019; Gaillochet *et al.*, 2022).

(b) Summary of genotyping by Sanger sequencing. One to 11 2-FA resistant individuals were genotyped for six different vector combinations (pCas12a, TaLbCas12a or UmLbCas12a-9i fused to YFP or not and targeting APT-1 or APT-2. Number of 2-FA resistant individuals is visualized, including the number of individuals with a mutation at the target site (mutant).
